# CLCC1 promotes membrane fusion during herpesvirus nuclear egress

**DOI:** 10.1101/2024.09.23.614151

**Authors:** Bing Dai, Lucas Polack, Adrian Sperl, Haley Dame, Tien Huynh, Chloe Deveney, Chanyoung Lee, John G. Doench, Ekaterina E. Heldwein

## Abstract

*Herpesvirales* are an ancient viral order that infects species from mollusks to humans for life. During infection, these viruses translocate their large capsids from the nucleus to the cytoplasm independently from the canonical route through the nuclear pore. Instead, capsids dock at the inner nuclear membrane and bud into the perinuclear space. These perinuclear enveloped virions fuse with the outer nuclear membrane releasing the capsids into the cytoplasm for maturation into infectious virions. The budding stage is mediated by virally encoded proteins. But the mediator of the subsequent fusion stage is unknown. Here, using a whole-genome CRISPR screen with herpes simplex virus 1, we identified CLCC1 as an essential host factor for the fusion stage of nuclear egress. Loss of CLCC1 results in a defect in nuclear egress, accumulation of capsid-containing perinuclear vesicles, and a drop in viral titers. In uninfected cells, loss of CLCC1 causes a defect in nuclear pore complex insertion. Viral homologs of CLCC1 are present in herpesviruses that infect mollusks and fish. Our findings uncover an ancient cellular membrane fusion mechanism important for the fundamental cellular process of nuclear envelope morphogenesis that herpesviruses hijack for capsid transport.

## INTRODUCTION

*Herpesvirales* are large, enveloped viruses that infect much of the animal kingdom. The order is divided into three families: *Malacoherpesviridae* infect mollusks, *Alloherpesviridae* infect fish and amphibians, and *Herpesviridae*, commonly known as herpesviruses, infect mammals, birds, and reptiles, and cause lifelong infections in most of the world’s population. The family *Herpesviridae* is further subdivided into three subfamilies, *Alpha-, Beta-, and Gammaherpesvirinae*. Nine human herpesviruses from these three subfamilies cause diseases ranging from skin lesions to life-threatening eye ailments, encephalitis, cancer, and developmental abnormalities. No cures exist, and prophylactic and therapeutic options are limited.

Despite substantial sequence divergence across *Herpesvirales,* key replication steps are conserved, one being nuclear egress. Herpesviruses replicate their double-stranded DNA genomes and package them into capsids within the nucleus. Genome-containing capsids are then exported into the cytoplasm for maturation into infectious virions. Many eukaryotic viruses that replicate their genomes within the nucleus, such as HIV, influenza, and papillomaviruses, escape this double-membraned organelle via the canonical export pathway through the nuclear pore complex (NPC) ^1^. But the ∼40-50-nm opening of the nuclear pore ^2^ is too small to accommodate the ∼125-nm capsids of herpesviruses. So, *Herpesvirales,* instead, use a different, more complex nuclear export route termed nuclear egress ^3^. First, capsids dock and bud at the inner nuclear membrane (INM), forming perinuclear enveloped virions (PEVs) (budding, or envelopment). PEVs then fuse their temporary envelopes with the outer nuclear membrane (ONM), releasing unenveloped capsids into the cytoplasm (fusion, or de-envelopment).

The budding stage is mediated by the virally encoded UL31 and UL34 proteins that form the heterodimeric nuclear egress complex (NEC). The NEC has an intrinsic ability to deform and bud membranes by forming a hexagonal membrane-bound scaffold ^4^. UL31 and UL34 are essential for nuclear egress across *Herpesviridae* ^5-11^, and their homologs are found in all family members^3^. By contrast, proteins that facilitate the fusion stage have not yet been identified. Viral entry glycoproteins gB and gH have been proposed to mediate the fusion stage in HSV-1, but their individual knockouts have mild, if any, phenotypes ^12^.

This raised the possibility that herpesviruses might use the host fusion machinery during the fusion stage. Host processes that involve nuclear envelope membrane fusion include the NPC insertion during interphase and nuclear budding used to export large RNPs or misfolded proteins [reviewed in ^13,14^]. However, the fusogen that mediates these processes has not yet been identified. If herpesviruses hijacked this process during nuclear egress, identifying host factors involved in herpesvirus nuclear egress could, potentially, reveal the fusogen mediating fusion of the nuclear envelope.

Towards this goal, here we developed a quantitative flow-cytometry-based assay to measure capsid nuclear egress in the prototypical herpes simplex virus 1 (HSV-1) and used it in conjunction with a whole-genome CRISPR-Cas9 screen. The top hit in our screen was CLCC1, an ER chloride channel ^15,16^. We show that CLCC1 is essential for the fusion stage of HSV-1 nuclear egress. Loss of CLCC1 resulted in a defect in HSV-1 nuclear egress, accumulation of capsid-containing PEVs, and a drop in viral titers. In uninfected cells, loss of CLCC1 induced a phenotype associated with a defect in NPC insertion. Loss of CLCC1 also decreased viral titers in the closely related herpes simplex virus 2 (HSV-2) and pseudorabies virus (PRV). Expression of the wild-type CLCC1 *in trans* rescued these defects.

Intriguingly, homologs of CLCC1 are encoded in the genomes of *Malacoherpesviridae* and *Alloherpesviridae*, which infect mollusks and fish, respectively. This suggests that CLCC1 function may be important for herpesviral replication across the entire order *Herpesvirales* and raises questions about their evolutionary origins.

Collectively, our results show that CLCC1 facilitates membrane fusion during NPC insertion and during capsid nuclear egress in herpesviruses. Our findings link nuclear envelope fusion in herpesviruses and the host, illuminating an ancient cellular membrane fusion mechanism crucial for nuclear envelope morphogenesis that has been co-opted by herpesviruses.

## RESULTS

### CRISPR-Cas9 screens of nuclear egress identify CLCC1 as a top positive regulator

Herpesvirus nuclear egress is typically quantified by visualizing infected cells by transmission electron microscopy (TEM) and counting capsids in the nucleus, PNS, and cytoplasm. However, this assay is labor-intensive and can only be performed on a small scale. To increase the scale and throughput, we developed a flow-cytometry-based nuclear egress assay that combines partial membrane permeabilization with capsid-specific immunostaining to detect cytoplasmic capsids. Infected HeLa cells were first treated with digitonin, a mild detergent that permeabilizes the plasma membrane but not the nuclear envelope. Permeabilized cells were then stained with a primary antibody, 8F5, that binds HSV-1 major capsid protein VP5 on capsids but does not bind free VP5 ^17^ and an Alexa 488-conjugated secondary antibody. To ensure that analyzed cells were infected, we used an HSV-1 F strain GS3217 encoding an NLS-tdTomato transgene expressed from an immediate-early (IE) promoter ^18^. We monitored two fluorescence channels: tdTomato, for the detection of HSV-1 infection, and Alexa-488, for the detection of capsids. Detection of double-positive tdTomato+/Alexa488+ cells served as a readout for nuclear egress (**Fig 1a, Extended Data Fig. 1a, 1c)**. The HSV-1 mutant lacking UL34, an NEC component, served as a negative control (**Fig 1a, Extended Data Fig. 1b, 1c)**. In the WT HSV-1, ∼90% cells were tdTomato+/Alexa488+ (**Fig 1a)** whereas in the UL34-null mutant, only ∼6 % of cells were tdTomato+/Alexa488+ (**Fig 1b)**. To rule out defects in capsid production, infected cells were fully permeabilized with Triton-X100, which permeabilizes both the plasma membrane and the nuclear envelope, as a control. In the fully permeabilized WT or UL34-null HSV-1, ∼95% of cells are tdTomato+/Alexa488+ (**Extended Data Fig. 1a-1c)**.

**Figure 1.**
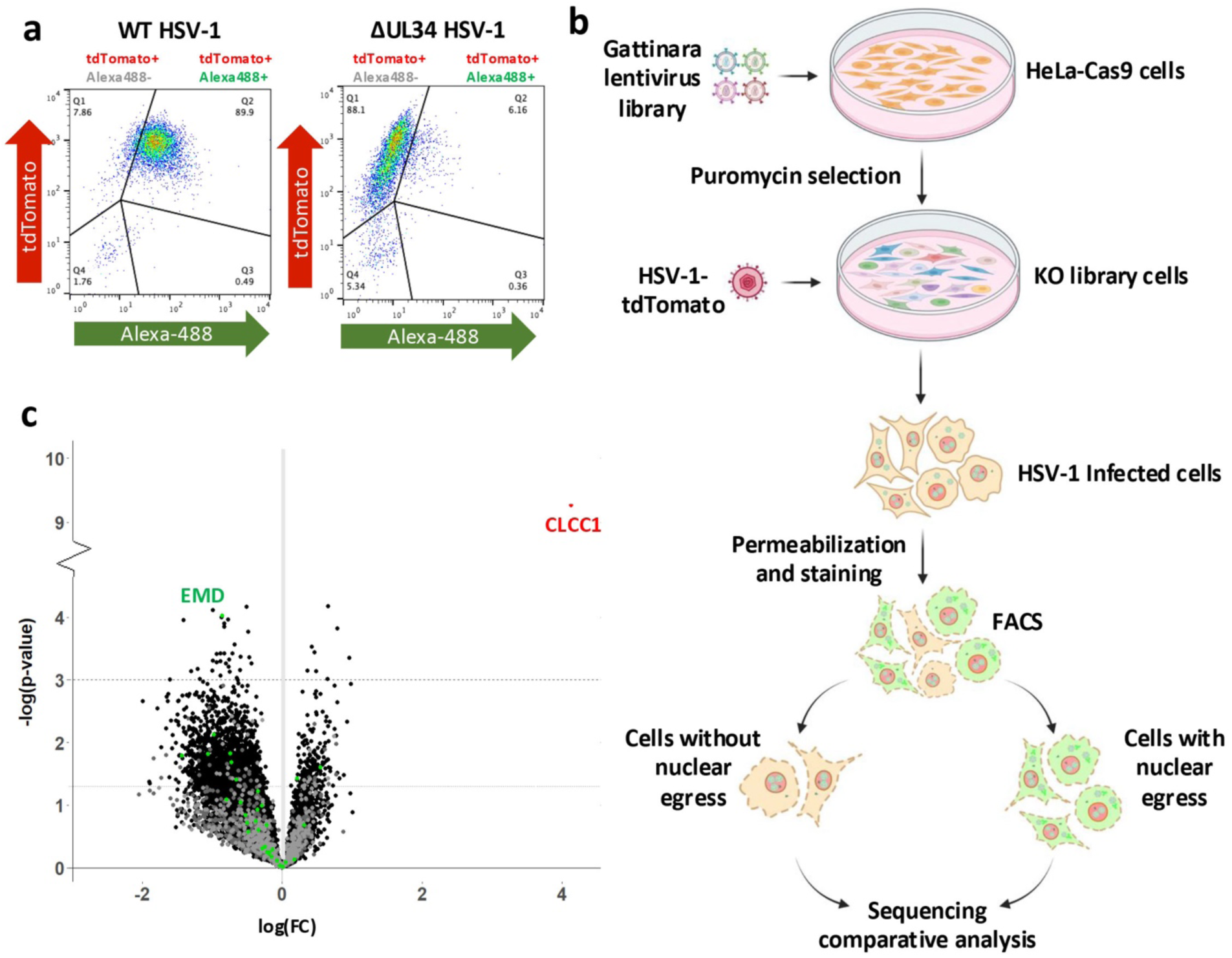
CLCC1 emerged as a top positive regulator of HSV-1 nuclear egress in a genome-wide CRISPR-Cas9 screen. **a)** The flow-cytometry-based nuclear egress assay separates HSV-1 infected HeLa cells with vs. without nuclear egress based on two fluorescent signals: tdTomato (red, HSV-1 infection) and Alexa-488 (green, presence of cytoplasmic capsids). HeLa cells infected with HSV-1 encoding tdTomato, were partially permeabilized 24 hpi and stained with an anti-capsid mAb and an Alexa488-conjugated secondary mAb. Cells in the tdTomato+/Alexa488+ quadrant (Q2) are infected and have cytoplasmic capsids, indicating nuclear egress. Cells in the tdTomato+/Alexa488-quadrant (Q1) are infected but do not have cytoplasmic capsids, indicating no nuclear egress. Left: ∼90% cells infected with WT HSV-1 are tdTomato+/Alexa488+. Right: only ∼6% cells infected with HSV-1 ΔUL34 virus, which has a defect in nuclear egress, are tdTomato+/Alexa488+. **b)** Schematic of the genome-wide CRISPR screen. HeLa-Cas9 cells were transduced with Gattinara library lentivirus, containing ∼40,000 sgRNAs, with 2 sgRNAs/gene, and after selection with puromycin, were infected with HSV-1 encoding tdTomato. 24 hpi, partially permeabilized cells were stained with a capsid-specific antibody and sorted by flow cytometry. **c)** Volcano plot of the screen results. Each dot represents a specific gene. The x-axis shows the fold change (FC) of sgRNAs, plotted as log(FC). Genes with log(FC) values >0 or <0 are candidate positive or negative regulators, respectively. The y-axis shows the significance score plotted as -log(p-value). The dotted line at y = 3 is the threshold for p-value < 0.001, indicating high confidence candidates. The dashed line at y = 1.3 is the threshold for p-value < 0.05. Red: top hit, CLCC1. Green: genes known to contribute to HSV-1 nuclear egress. High-confidence hit, EMD, is labelled. Gray: control sgRNAs (targeted, non-site, or intergenic sites).

To identify host factors involved in nuclear egress, we performed a CRISPR-Cas9 screen in HeLa cells. A Cas9-expressing HeLa cell line was transduced with the Gattinara sgRNA library composed of ∼40,000 sgRNAs targeting the whole human genome, with two sgRNAs per gene. Transduced cells were infected with WT HSV-1, fixed, partially permeabilized, stained, and sorted by fluorescence-activated cell sorting (FACS). Two cell populations – with or without nuclear egress – were collected. The tdTomato+/Alexa488-cell population (no nuclear egress) was analyzed for potential hits. The tdTomato+/Alexa488+ cell population (nuclear egress) was used as a control.

Genomic DNA was isolated from the sorted populations and sequenced **(Fig 1b).** Two independent Gattinara library transductions were done (2 biological replicates), each with three independent HSV-1 infections (3 technical replicates), with an R^2^ of 0.0012 (**Supplementary Fig 1**). The screen yielded 41 high-confidence candidate regulators with the p-value less than 0.001, including 9 positive regulators (decreased nuclear egress when the gene is depleted) (**Supplementary Table 1a**), and 32 negative regulators (increased nuclear egress when the gene is depleted) (**Supplementary Table 1b**). Among these, one host factor, EMD, was previously reported as contributing to HSV-1 nuclear egress, emerged as a negative regulator in our screen (**Fig 1c, Supplementary Table 1b)**. EMD encodes Emerin, a component of the nuclear lamina that helps maintain nuclear envelope integrity. Phosphorylation of emerin during HSV-1 infection promotes lamina disruption, which facilitates capsid nuclear egress ^19,20^. The presence of a host factor with known contributions to HSV-1 nuclear egress among the screen hits supports the validity of our screening approach. The rest of the previously reported host factors [reviewed in ^21^] had p-values higher than 0.001 (**Supplementary Table 2**).

The strongest positive regulator and the only hit present in all the replicates and with both sgRNAs was CLIC-like 1 chloride channel (CLCC1) (**Fig 1c, Supplementary Fig 1)**. CLCC1 is an ER channel ^15,16^ that mediates chloride efflux from the ER to neutralize charge imbalance caused by calcium release ^15^. CLCC1 plays a role in the ER stress and the unfolded protein response (UPR) ^15,22,23^. In humans, mutations in CLCC1 are associated with Amyotrophic Lateral Sclerosis (ALS) ^15^ and autosomal recessive retinitis pigmentosa ^24^. How CLCC1 could be involved in nuclear egress was unclear.

### Loss of CLCC1 causes a defect in HSV-1 capsid nuclear egress

To validate the defect in nuclear egress due to CLCC1 depletion, we generated CLCC1 knockout HeLa cell lines with two CLCC1-targeting sgRNAs from the Brunello library (sgRNAs CLCC1-3 and CLCC1-6), which were different from the ones used in the primary screens. From the heterogenous (bulk) pools of cells transduced with individual sgRNAs (cko3_bulk and cko6_bulk) (**Extended Data Fig 2a**), four single-cell clones (cko3_2, cko3_4, cko6_1, and cko6_2) were selected (**Fig 2a, Extended Data Fig 2a**). As a negative control, a HeLa cell line was transduced with an sgRNA targeting an intergenic region (Int_bulk) (**Extended Data Fig 2a**), and two single clones (Int_3 and Int_4) were selected (**Fig 2a, Extended Data Fig 2a)**. All bulk and single-cell CLCC1-KO cell lines had defects in HSV-1 nuclear egress as measured by the flow cytometry assay **(Fig 2a, Extended Data Fig. 2a)** and confirmed by confocal microscopy (**Extended Data Fig. 3**). Three single-cell CLCC1-KO clones, cko3_4, cko6_1, and cko6_2, showed strong defects in nuclear egress, <20%, comparable to that of the control UL34-null HSV-1 mutant, whereas cko3_2 had a more modest defect, ∼40% **(Fig 2a, Extended Data Fig. 2a).** These data validated CLCC1 as a host factor required for HSV-1 capsid nuclear egress. Two of the single-cell CLCC1-KO clones, cko3_4 and cko6_1, were chosen for further characterization. One single clone targeting an intergenic region, Int_4, was chosen as a negative control.

**Figure 2.**
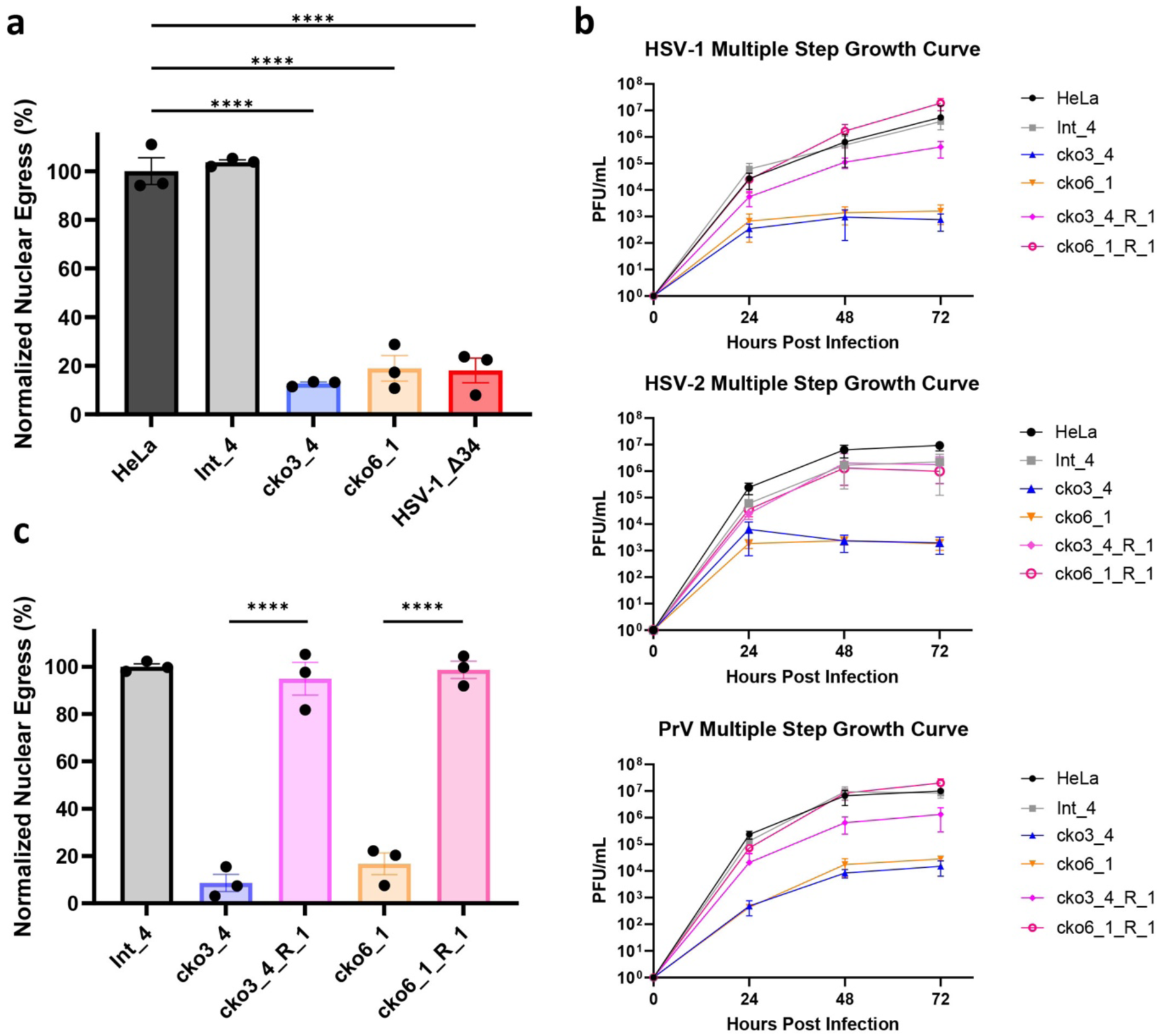
CLCC1 is essential for HSV-1 nuclear egress and HSV-1, HSV-2, and PrV replication. **a)** Depletion of CLCC1 causes a defect in nuclear egress, measured by the flow cytometry nuclear egress assay. Single-clone CLCC1-KO (cko3_4 and cko6_1) or two control, HeLa and intergenic site targeting (Int_4) cell lines were infected with WT HSV-1 at an MOI of 5. As a positive control, HeLa cells were infected with HSV-1 ΔUL34 mutant virus, defective in nuclear egress, at an MOI of 10. Nuclear egress was measured at 24 hpi and normalized to HSV-1-infected HeLa cells. Each experiment had three biological replicates, each containing two technical replicates. Each data point represents a biological replicate. Bars represent mean values, and the error bars represent SEM. P < 0.0001 = ****. Significance was calculated using one-way ANOVA, with multiple comparisons. **b)** Multiple-step growth curves for HSV-1 on single-clone CLCC1-KO (cko3_4 and cko6_1), single-clone CLCC1-R (cko3_4_R_1 and cko6_1_R_1), or two control, HeLa and Int_4, cell lines. Cells were infected with HSV-1 at MOI of 0.1, and supernatants were titrated in Vero cells using plaque assay. The y-intercept is set to 10^0^ at time 0 for visual clarity. Each symbol shows the mean of three biological replicates, each containing two technical replicates, and bars show SEM. **c)** Expression of CLCC1 in trans rescues the defect in nuclear egress, measured by the flow cytometry nuclear egress assay as in **a).** Single-cell CLCC1-KO (cko3_4 and cko6_1), single-cell CLCC1-R (cko3_4_R_1 and cko6_1_R_1), or control Int_4 cell lines were infected with HSV-1 at an MOI of 5. Nuclear egress was measured at 24 hpi and normalized to Int_4 signal. Each data point represents a biological replicate. Each experiment had three biological replicates, each containing two technical replicates. Each data point represents a biological replicate. Bars represent mean values, and the error bars represent SEM. P < 0.0001 = ****. Significance was calculated using one-way ANOVA, with multiple comparisons.

### Loss of CLCC1 causes defects in viral replication in HSV-1 and two related *Alphaherpesvirinae*

To examine defects on viral replication due to loss of CLCC1, we measured HSV-1 titers using multiple-step growth curves. HSV-1 replication in either cko3_4 or cko6_1 cell lines resulted in a ∼1000-fold drop in titer **(Fig 2b)**. The strong defect in virion production due to the loss of CLCC1 is consistent with the strong defect in nuclear egress, which is an essential step in virion morphogenesis. To determine if CLCC1 were important for replication in other members of the *Alphaherpesvirinae* subfamily of the family *Herpesviridae*, we tested HSV-2 and pseudorabies virus (PRV). Replication of both HSV-2 and PRV in both CLCC1-KO cell lines resulted in ∼1000-fold drop in titer (**Fig 2b)**. Thus, CLCC1 is required for nuclear egress across several members of the *Alphaherpesvirinae* subfamily.

### The defects in nuclear egress and viral replication due to loss of CLCC1 are rescued by expression of CLCC1 *in trans*

To confirm that the defects in nuclear egress and viral replication in CLCC1-KO cells were specific to the loss of CLCC1, we performed a rescue experiment by expressing CLCC1 *in trans*. To do so, we generated the CRISPR-resistant (CR) gene variant of CLCC1 (CLCC1-CR), in which silent mutations were introduced to destroy the target sites for 6 CLCC1 sgRNAs (2 from Gattinara and 4 from Brunello libraries). Transient or stable overexpression of CLCC1-CR under control of a strong promoter reduced HSV-1 nuclear egress in Int_4 cells and poorly rescued the nuclear egress defect in CLCC1-KO cells (**Extended Data Fig 4a, 4b)**. Therefore, we stably expressed CLCC1-CR under control of a weak promoter in cko3_4, cko6_1, and Int_4 cell lines (**Extended Data Fig 4b)**. From the bulk rescue pools cko3_4_R_bulk and cko6_1_R_bulk (**Extended Data Fig 4a-4c)**, several single-cell CLCC1 rescue (CLCC1-R) clones were selected (**Extended Data Fig 4c, 4d)**. Partial rescue of the nuclear egress defect due to loss of CLCC1, between 30-80%, was observed in bulk and some single-cell CLCC1-R clones **(Extended Data Fig 4c).** Full rescue of the nuclear egress defect (>90%) was observed in single-cell CLCC1-R clones cko3_4_R_1 and cko6_1_R_1 **(Fig 2c, Extended Data Fig 4c).** Replication of HSV-1, HSV-2, and PRV was also rescued nearly to the WT levels **(Fig 2b).** These results confirmed the importance of CLCC1 in both HSV-1 nuclear egress and viral replication across the *Alphaherpesvirinae* subfamily.

### The CLCC1 role in nuclear egress is unrelated to ER stress and UPR

CLCC1 has been linked to ER stress and an unfolded protein response (UPR) ^15,22,23^. To evaluate the role of ER stress in the HSV-1 nuclear egress, we measured levels of BiP, a mediator of UPR and an ER stress marker ^25^. In uninfected HeLa or single-cell CLCC1-KO (cko6_1) cell lines, BiP levels were higher upon treatment with a chemical ER stress inducer dithiothreitol (DTT) (**Extended Data Fig 5a**). cko6_1 cells were also more sensitive to DTT than HeLa cells, judging by their lower viability (**Extended Data Fig 5b**). However, during HSV-1 infection, BiP levels were similarly low in the presence or absence of DTT, in both HeLa and CLCC1-KO cell lines (**Extended Data Fig 5a**). HSV-1 is known to suppress UPR ^26,27^. Importantly, HSV-1 nuclear egress is not inhibited by DTT in HeLa cells (**Extended Data Fig 5c**). Therefore, ER stress is unlikely to explain the HSV-1 nuclear egress defect due to the loss of CLCC1.

### PEVs accumulate in the perinuclear space of HSV-1-infected cells in the absence of CLCC1

To determine the stage in nuclear egress blocked in the absence of CLCC1, we examined single-clone CLCC1-KO (cko3_4 and cko6_1) and CLCC1-R (cko6_1_R_1) cell lines infected with HSV-1 by using transmission electron microscopy (TEM). HSV-1-infected Int_4 cell line was used as a control. In HSV-1-infected CLCC1-KO cell lines, PEVs accumulated in the PNS **(Fig 3a, 3b**), indicating a defect at the fusion stage of nuclear egress. By contrast, in the control Int_4 and CLCC1-R cell lines, only single PEVs were observed **(Fig 3a).** Thus, CLCC1 may facilitate the de-envelopment (fusion) stage of capsid nuclear egress.

**Figure 3.**
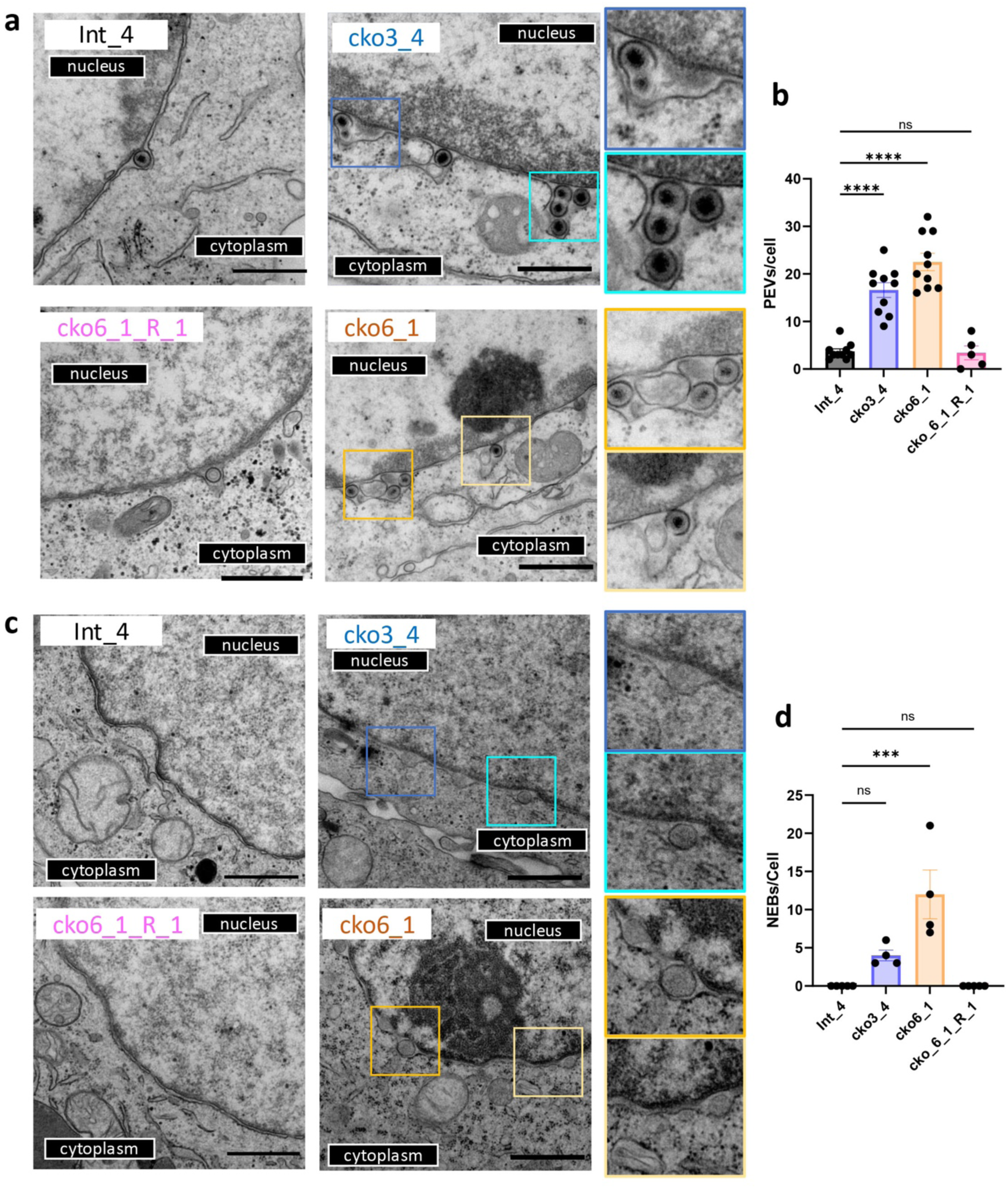
Depletion of CLCC1 causes accumulation of PEVs in HSV-1-infected cells and formation of nuclear enveloped blebs (NEBs) in uninfected cells. **a,c)** TEM images of Int_4, CLCC1-KO (cko3_4 and cko6_1), and CLCC1-R (cko6_1_R_1) cell-lines either infected with HSV-1 at an MOI of 5 **(a)** or uninfected **(c).** Scale bar = 800 nm. Zoomed-in views of features of interest are shown on the right. **b, d)** Quantification of PEVs in infected cells **(b)** and NEBs in uninfected cells **(d).** In **(b),** data were combined from two biological replicates. Each dot represents the number of events in a single cell. Bars represent mean values, and the error bars represent SEM. P < 0.001 = ***; P < 0.0001 = ****. Significance was calculated using one-way ANOVA, with multiple comparisons.

### Loss of CLCC1 in uninfected cells causes formation of blebs due to a defect in NPC insertion

As a control, we examined uninfected single-clone CLCC1-KO (cko3_4 and cko6_1) cell lines by TEM. Surprisingly, we observed vesicles, or blebs, in the PNS **(Fig 3c, 3d).** Unlike in HSV-1-infected CLCC1-KO cell lines, where multiple PEVs accumulated, these blebs did not accumulate. Instead, the blebs were distributed along the nuclear envelope, giving it a bead-like appearance. No blebs were found in the control Int_4 and CLCC1-R cell lines **(Fig 3c, 3d).** Up-close examination revealed that some blebs had necks and appeared connected to the INM.

Morphologically similar blebs have been observed in cells depleted of the Torsin ATPase or its cofactors LAP1 and LULL1 ^28,29^. This phenotype was attributed to a defect in NPC insertion during interphase nuclear pore biogenesis caused by a defect in the fusion of the inner and outer nuclear membranes ^28^. Myeloid leukemia factor 2 (MLF2) has been identified as a component of the bleb lumen ^28^. To test for the presence of MLF2 in blebs formed in cells lacking CLCC1, we overexpressed an MLF2 construct fused to a GFP reporter. We found that MLF2 localized to blebs along the nuclear envelope in CLCC1-KO cell lines (cko3_4 and cko6_1) but not in the control HeLa and Int_4 cells or the CLCC1-R cell lines (cko3_4_R_1 and cko6_1_R_1) (**Extended Data Fig 6**). Thus, loss of CLCC1 recapitulated the nuclear blebbing phenotype previously observed in cells depleted of Torsin and attributed to a defect in NPC insertion. We conclude that the CLCC1 is required not only for HSV-1 nuclear egress but also for nuclear pore biogenesis during interphase.

### Members of the order *Herpesvirales* encode CLCC1 homologs

There are >1000 CLCC1 homologs across the animal kingdom. Unexpectedly, we discovered viral homologs of CLCC1 (vCLCC1) in four *Malacoherpesviridae*, Oyster herpesvirus 1 (OsHV-1), Malacoherpesvirus 1 (MLHV1), Abalone herpesvirus (AbHV), and Chlamys acute necrobiotic virus (CanV); and four *Alloherpesviridae*, Ictalurid herpesvirus 1 (IcHV-1), Anguilid herpesvirus 1 (AngHV-1), Black bullhead herpesvirus (BbHV), and Silurid herpesvirus 1 (SHV-1) (**Supplementary Table 3**). vCLCC1s and cellular CLCC1s (cCLCC1) are predicted to have three transmembrane (TM) helices with the same topology, with N and C termini predicted to face the ER and the cytoplasm, respectively (N_ER_-TM1-TM2-TM3-C_cyto_) (**Fig 4a, 4b**). The ∼180 amino acid “core” region of CLCC1 from TM1 to TM3 is highly conserved across all homologs, with very similar predicted folds (**Fig 4b, 4c, and Supplementary Fig 2**). By contrast, the N termini adopt different folds, and the C termini are largely disordered across all homologs (**Fig 4b**). The vCLCC1s are also shorter than cCLCC1s by ∼150 amino acid residues due to shorter N and C termini (**Fig 4a and Supplementary Fig 2**).

**Figure 4.**
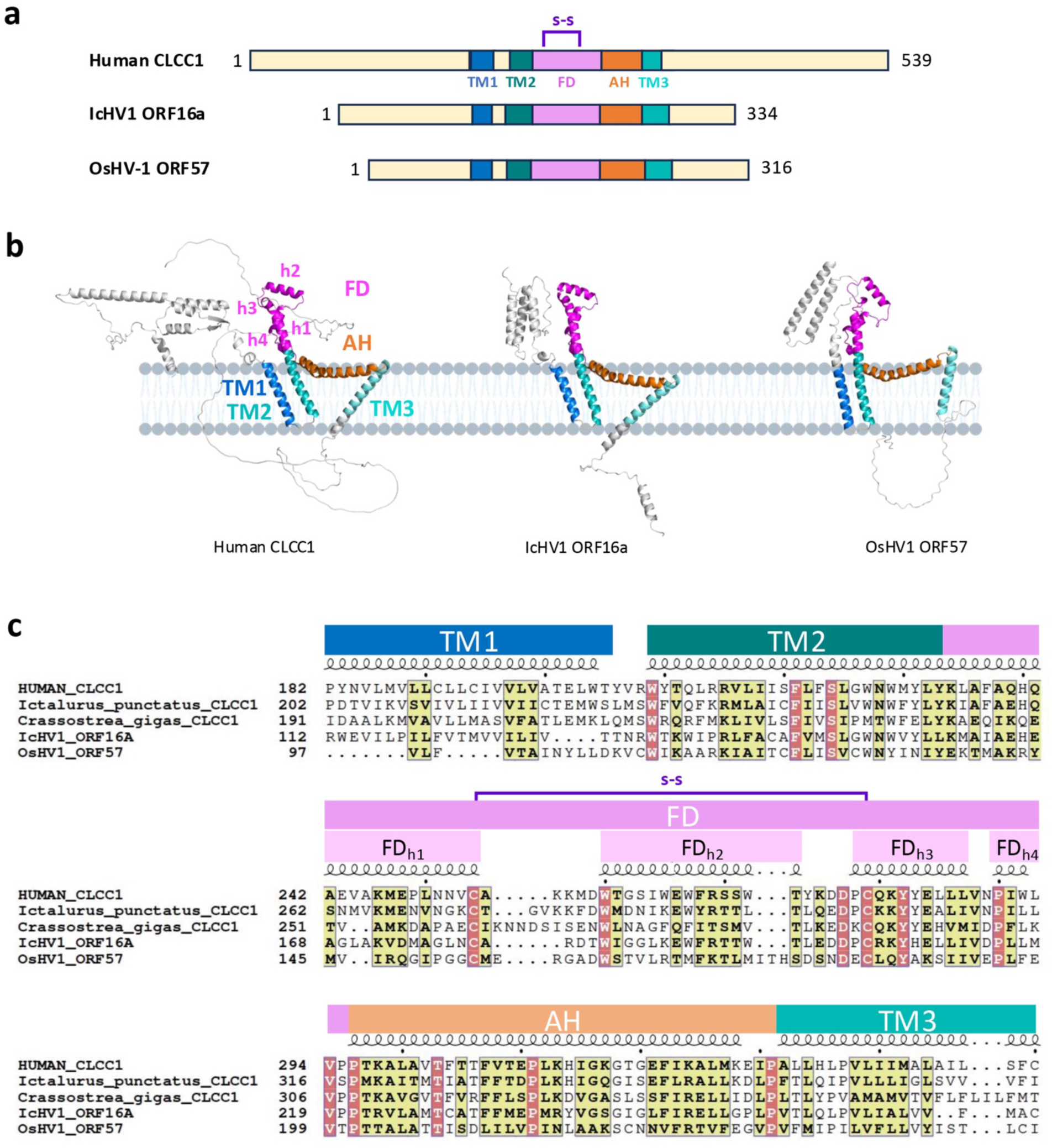
Herpesviral and cellular CLCC1 homologs share sequence and structural similarity. **a)** Domain architecture of human CLCC1 and two herpesviral CLCC1 homologs, IcHV1 ORF16a and OsHV-1 ORF57. Structural elements and domains were assigned based on structural predictions and secondary structure assignments and colored as follows: transmembrane helix 1 (TM1, blue), TM2 (deep teal), fist domain (FD, magenta), amphipathic helix (AH, orange), and TM3 (teal). S-S = predicted disulfide bond (purple). Transmembrane domains were predicted by TMHMM 2.0 (https://services.healthtech.dtu.dk/services/TMHMM-2.0/). **b)** Ribbon diagrams of AlphaFold3 models of human CLCC1, IcHV1 ORF16a, and OsHV-1 ORF57. Structural models were generated using the AlphaFold 3.0 online server (https://golgi.sandbox.google.com/) and displayed using Pymol. Structural elements and domains are colored as in **(a)** and labelled. **c)** Multiple sequence alignment of human CLCC1, its herpesviral homologs IcHV1 ORF16a and OsHV-1 ORF57, and CLCC1 homologs from the respective hosts, Ictalurus punctatus (Channel catfish) and Crassostrea gigas (Pacific oyster). Similar residues are highlighted in yellow. Identical residues are highlighted in red. Structural elements and domains are colored and labelled as in **(a)** and **(b)**. Sequence alignment was generated using Clustal Omega^44^ and rendered using ESPript 3.0^45^ (https://espript.ibcp.fr).

AlphaFold3 ^30^ predicts similar folds for the cCLCC1 homologs (e.g., human CLCC1) and vCLCC1 homologs from *Alloherpesviridae* (e.g., IcHV1 ORF16a) and *Malacoherpesviridae* (e.g., OsHV-1 ORF57) (**Fig 4b**). These folds do not resemble any known structures ^31^. TM1 and TM2 are adjacent and antiparallel whereas TM3 is separate and tilted in respect to TM1/TM2 (**Fig 4b**). The tilt of TM3 is greater in human CLCC1 and IcHV1 ORF16a than OsHV-1 ORF57 **(Fig 4b)**. TM2 is followed by a fist domain (FD), composed of 4 helices of variable length, FD_h1_-FD_h4_. Helix FD_h1_, a continuation of TM2, is followed by an amphipathic helix FD_h2_ (the knuckle part of the fist) that is oriented perpendicular to helix FD_h1_ and juts outward from the membrane. Short helices FD_h3_ and FD_h4_ run antiparallel to helix FD_h1_. A highly conserved disulfide between the C terminus of FD_h1_ and the N terminus of FD_h3_ stabilizes the fold (**Fig 4a, 4c, and Extended Data Fig 7**). The FD is followed by a long, bow-shaped amphipathic helix (AH) flanked by invariant prolines (**Fig 4b and Extended Data Fig 7)**. Another invariant proline in the middle of AH gives it its bow shape **(Fig 4b and Extended Data Fig 7)**. AH is followed by TM3. TM1/TM2, AH, and TM3 form a triangular shape (**Fig 4b**). The tilted orientations of the TMs place AH in a position to interact peripherally with the membrane (**Fig 4b**).

The TM1-TM2-FD-AH-TM3 module is conserved in sequence and predicted secondary and tertiary structure (**Fig 4**). It also contains 10 residues that are invariant across 8 representative animal and 4 herpesviral homologs (**Extended Data Fig 7 and Supplementary Fig 2)**. Some of these, e.g., 4 prolines and 2 cysteines, are likely structurally important whereas others are likely functionally important.

### Highly conserved CLCC1 residue and residues involved in chloride channel activity are important for HSV-1 nuclear egress

To help define the mechanistic role of CLCC1 in HSV-1 nuclear egress, we tested the ability of CLCC1 mutants to rescue the nuclear egress defect caused by the loss of CLCC1. A prior study using human and mouse CLCC1 ^15^ reported several mutations that altered its channel function *in vitro* and *in vivo* (**Fig 5a, 5b**). D25E and D181R, which target a putative Ca^2+^-binding site, make CLCC1 less sensitive to Ca^2+^ inhibition and reduce Ca^2+^ binding *in vitro* ^15^. Additionally, D25E is associated with autosomal recessive retinitis pigmentosa ^24^. S263R and W267R reduce channel conductivity *in vitro* and are associated with ALS ^15^. K298E mutation reduces channel potentiation by PIP2 ^15^. All these mutations target ER-facing residues. To probe the role of an invariant residue, we mutated D277 to an arginine, reversing its charge. D277 was chosen because it is located within an ER-facing segment, FD, just like known functionally important residues described above (**Fig 5a, 5b**). As controls, we generated double mutants D152R/D153R and E175R/D176R that did not affect channel conductivity *in vitro* ^15^.

**Figure 5.**
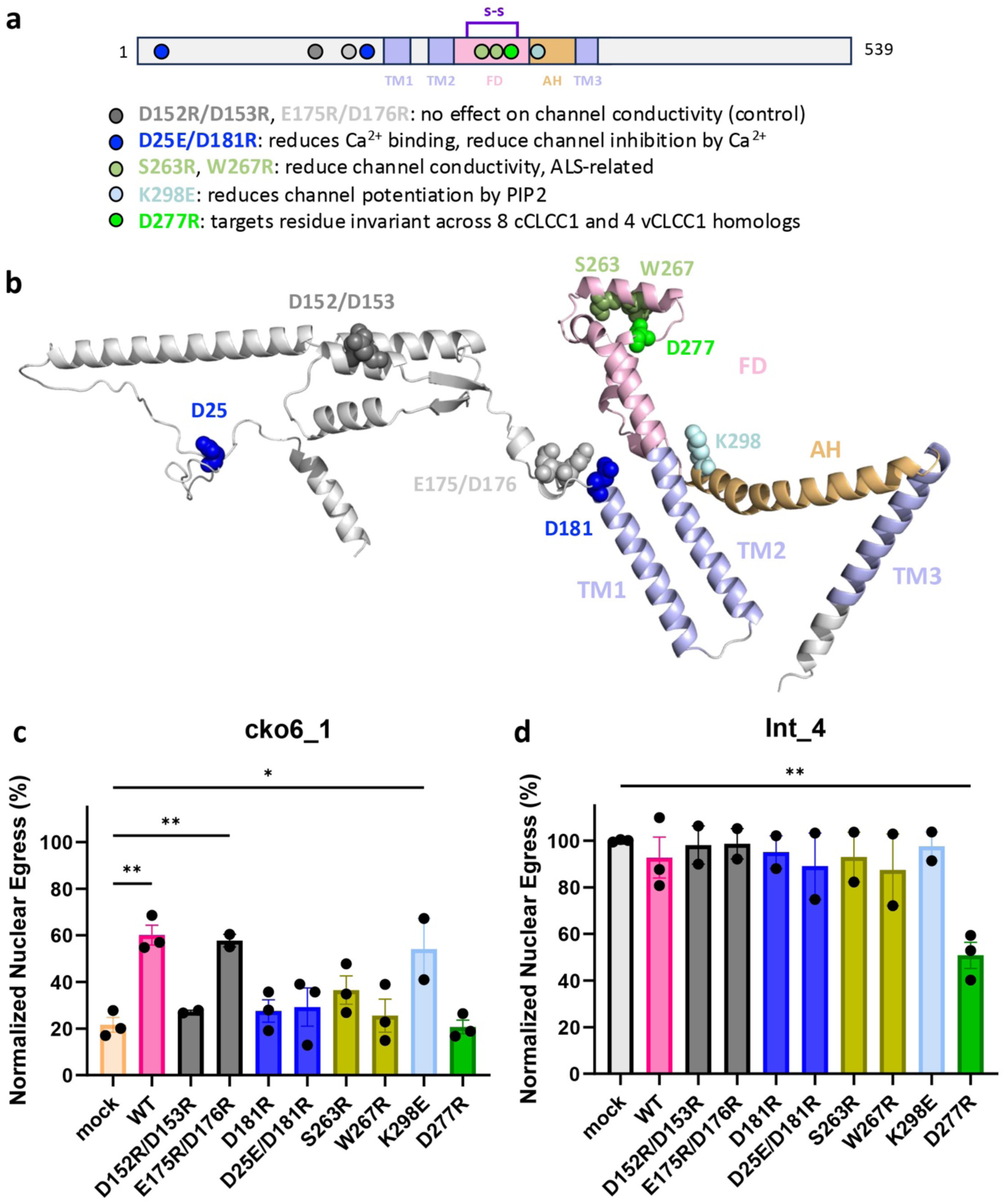
A highly conserved CLCC1 residue and residues involved in chloride channel activity are important for HSV-1 nuclear egress. **A)** Schematic representation of the locations of CLCC1 mutations and their functions (if known). Structural elements and domains were assigned as in Fig 4 and colored as follows: TM1/TM2/TM3 (light blue), FD (light pink), and AH (light orange). S-S = predicted disulfide bond (purple). Approximate locations of mutated residues are shown as dots. **b)** A ribbon diagram of an AlphaFold3 model of human CLCC1. Structural elements and domains are colored as in **(a)** and labelled. Mutated residues are shown in sphere representation and colored as in **a)**. Residues 365-539 were removed, for clarity. **c, d)** Several CLCC1 mutants rescue a defect in nuclear egress due to CLCC1 depletion whereas others do not, as measured by the flow cytometry nuclear egress assay. Single-clone CLCC1-KO (cko6_1) **(c)** or control Int_4 cell line **(d)** were either mock-transduced (Mock) or transduced with lentiviral constructs encoding WT CLCC1 or mutants D152R/D153R, E175R/D176R, D181R, D25E/D181R, S263R, W267R, D277R, K298E in the CLCC1-CR background. Bulk pools were infected with WT HSV-1 at an MOI of 5. Nuclear egress was measured at 24 hpi and normalized to the Int_4 mock. Each experiment had at least two biological replicates, each containing two technical replicates. Each data point represents a biological replicate. Bars represent mean values, and the error bars represent SEM. P < 0.01 = **, P < 0.05 = *. Significance was calculated using one-way ANOVA, with multiple comparisons. The color scheme is as in **(a)** and **(b).**

We introduced mutations D152R/D153R, E175R/D176R, D181R, D25E/D181R, S263R, W267R, K298E, and D277R into the CLCC1-CR gene. To perform the CLCC1 rescue experiment, WT CLCC1 or CLCC1 mutants were expressed in trans in the control cell line (Int_4) or CLCC1-KO cell line (cko6_1) under the control of a weak promoter. Expression of E175R/D176R or K298E mutants partially rescued the defect in HSV-1 nuclear egress caused by the loss of CLCC1 to ∼50%, similarly to the WT CLCC1 (**Fig 5c**). Thus, mutations E175R/D176R and K298E do not appear to impair nuclear egress. By contrast, expression of CLCC1 mutants that reduce channel conductivity *in vitro* (S263R and W267R) or reduce Ca^2+^ binding (D181R, D25E/D181R) did not rescue the HSV-1 nuclear egress defect (**Fig 5c**). Expression of the D277R mutant, which targets an invariant residue, or the control mutant D152R/D153R also did not rescue the nuclear egress defect (**Fig 5c**). Thus, mutations D152R/D153R, D181R, D25E/D181R, S263R, W267R, and D277R mutations impair nuclear egress. We hypothesize that chloride channel activity and Ca^2+^ binding might be important for the CLCC1 function in nuclear egress. The invariant residue D277 is also important for the CLCC1 function in HSV-1 nuclear egress. The expression of D277R in Int_4 cells, which have endogenous CLCC1, reduced nuclear egress, acting in a dominant-negative manner (**Fig 5d**). If so, D277 could potentially mediate CLCC1 oligomerization. The roles of D152 and D153 are yet unclear, but these residues are located on a predicted helix within the N-terminal ER-facing segment (**Fig 5b**) and could mediate protein-protein interactions.

We do not know how mutations in CLCC1 affect its cellular levels because we could not detect either the WT or the mutant CLCC1 proteins in bulk rescue experiments by Western Blot. Presumably, CLCC1 expression levels are below the detection limit of the Western Blot assay due to the use of a weak promoter. Indeed, WT CLCC1 was detected in single-cell clones isolated from the same bulk rescue pools (**Extended Data Fig 4d**). Additionally, all mutants tested here (except for D277R) were successfully expressed and purified previously ^15^, making poor expression or misfolding in our experiments unlikely. Finally, the D277R mutant reduces HSV-1 nuclear egress in cells expressing endogenous CLCC1, which suggests that it is expressed.

## DISCUSSION

### CLCC1 facilitates the fusion of the nuclear envelope in herpesvirus-infected and uninfected cells

Nuclear egress is an essential stage in replication conserved across all herpesviruses. For decades, this process was thought to be specific to herpesviruses until the discovery that Drosophila uses a topologically similar mechanism to export large mRNA/protein complexes during embryonic development ^32,33^. This non-canonical nuclear export pathway, referred to as nuclear envelope budding (NEB) among others, has also been proposed to export protein aggregates ^34^ as a response to stress ^35^. A similar nuclear blebbing (NB) phenotype was observed in cells depleted of the Torsin ATPase ^28,29^. This phenotype was attributed to a defect in NPC insertion during interphase nuclear pore biogenesis caused by a defect in the fusion of the inner and outer nuclear membranes ^28^. The relationship between NEB from NB is yet unclear. Nonetheless, the NEB/NB-like phenotypes have been reported in organisms spanning the range from yeast to sea urchins to mammals [reviewed in ^13,14^] as early as 1965 ^36^.

Despite morphological similarities, it was unclear whether herpesvirus nuclear egress and NEB/NB in eukaryotes shared any mechanistic similarities. In herpesviruses, the budding stage of nuclear egress is mediated by virally encoded UL31 and UL34 proteins [reviewed in ^3^] that have no known homologs outside of herpesviruses. Conversely, Torsin ATPase is essential for the budding stage of NEB in Drosophila ^32^ and NPC insertion ^28^ but dispensable for HSV-1 nuclear egress ^37^. Thus, herpesvirus nuclear egress and budding of the nuclear envelope in uninfected host cells use distinct budding mechanisms. But what factors facilitate membrane fusion of the nuclear envelope in either case have remained mysterious.

Here, by combining a whole-genome CRISPR-Cas9 screen with a custom nuclear egress assay, we identified CLCC1, an ER chloride channel, as a strong positive regulator of membrane fusion during HSV-1 nuclear egress. Loss of CLCC1 resulted in a defect in HSV-1 nuclear egress, accumulation of budded capsids in the perinuclear space, and a drop in viral titers. Loss of CLCC1 also reduced viral titers in the closely related herpes simplex virus 2 (HSV-2) and pseudorabies virus (PRV). Expression of the wild-type CLCC1 *in trans* rescued these defects. We also found that in uninfected cells, loss of CLCC1 caused a blebbing of the nuclear envelope associated with a defect in NPC insertion caused by a defect in the fusion of the inner and outer nuclear membranes. Our results show that CLCC1 not only facilitates membrane fusion during capsid nuclear egress in HSV-1 and, likely, closely related *Alphaherpesvirinae* HSV-2 and PRV but also facilitates NPC morphogenesis in the host. The nuclear egress process is found across the entire order *Herpesvirales*. Therefore, we propose that these viruses hijack the machinery that mediates nuclear envelope fusion during NPC insertion and nuclear budding for capsid nuclear egress, linking nuclear envelope fusion in herpesviruses and their animal hosts.

### The existence of viral CLCC1 homologs raises questions about their functions and evolutionary origins

We discovered viral homologs of CLCC1 in several members of the order *Herpesvirales*, four *Malacoherpesviridae* that infect mollusks (oysters, snails, abalone, and scallops) and four *Alloherpesviridae* that infect fish. Why these herpesviruses encode CLCC1 homologs is yet unclear given that their respective hosts encode their own CLCC1 homologs. For example, *Crassostrea gigas* (Pacific oyster) and *Ictalurus punctatus* (Channel Catfish), the respective hosts of OsHV-1 and IcHV-1, encode CLCC1 homologs (**Fig 4c, Supplementary Fig 2**). vCLCC1 homologs are shorter, however, and could have distinct functions. Importantly, the existence of vCLCC1s suggests that CLCC1 is important for herpesviral replication across the entire order *Herpesvirales*. More generally, it also raises questions about their evolutionary origins. We have not yet found any CLCC1 homologs in *Herpesviridae*, which infect mammals, birds, and reptiles. We note, however, that vCLCC1 homologs reported here were difficult to find using available homology search algorithms. More advanced types of homology search could identify additional homologs in *Malacoherpesviridae, Alloherpesviridae* and, possibly, *Herpesviridae*.

vCLCC1s and cCLCC1s have conserved ∼180 amino acid cores that are predicted to have very similar structural folds (**Fig 4b**) that do not resemble any known structures ^31^. The common fold consists of adjacent TM1 and TM2 that are separated from TM3 by a disulfide-stabilized helical FD and a long bow-shaped AH, TM1-TM2-FD-AH-TM3. It also contains 10 residues that are invariant across 8 representative cellular and 4 herpesviral CLCC1 homologs (**Supplementary Fig 2)**. Six of these are likely structurally important. The 2 cysteines, C254 and C279, are predicted to form a disulfide that likely stabilizes the FD (**Fig 4b and Extended Data Fig 7)**. Four prolines, P290, P296, P311, and P331 are located at the junctions of helices (FD_h3_-FD_h4_, FD_h4_-AH, AH-TM3) and in the middle of AH (**Fig 4b and Extended Data Fig 7**), consistent with their ability to disrupt helices or generate kinks, and likely stabilize the unusual fold of CLCC1. High conservation of these six residues suggests that the FD-AH-TM3 is an essential, structurally conserved element in the CLCC1 structure. The remaining four residues, W209, S224, D277, and Y282 likely have important functional roles. Indeed, D277, is important for HSV-1 nuclear egress because D277R mutant fails to rescue the nuclear egress defect caused by the loss of CLCC1 (**Fig 5**).

### How does CLCC1 promote membrane fusion of the nuclear envelope?

Purified CLCC1 has a chloride channel activity that is inhibited by Ca^2+^ ^15^. However, CLCC1 does not resemble its namesakes, the dimeric CLC channels ^38^, or any other ion channels in sequence or structural predictions. Native and recombinant CLCC1 form oligomers of unclear stoichiometry^15^. Ion channels typically form dimers, tetramers, or hexamers ^39^. But AlphaFold3 predictions of <10 CLCC1 copies generate oligomeric stacks (**Extended Data Fig 8**). Structural predictions of 10 or more copies form oligomeric rings with openings of increasing sizes all of which are too large for an ion channel (**Extended Data Fig 8**). Experimentally determined structures of CLCC1 homologs are needed to clarify its oligomeric state and function.

Mutations that reduce CLCC1 channel activity or make it less sensitive to Ca2+ inhibition affect CLCC1 function in herpesvirus nuclear egress (**Fig 5**). So, chloride channel function is likely important for herpesvirus nuclear egress. Although AlphaFold3 structural models do not pinpoint the location of the chloride-conducting pore within CLCC1, one side of TM2 is lined with hydrophilic residues (S220, S224, N228, and Y231), several of which are conserved and one, S224, invariant (**Extended Data Fig 7)**. Thus, TM2 could potentially participate in chloride transport across the channel.

How a chloride channel activity could facilitate membrane fusion is unclear. However, fusogenic activity of many membrane fusogens is controlled by pH and, in some cases, ions ^40^. Additionally, chloride is a major proton counterion. Therefore, CLCC1 could control the fusogenic activity of a yet unidentified fusogen by changing the pH or the osmotic environment of the perinuclear space.

Our CRISPR-Cas9 screens did not yield any strong positive regulators of nuclear egress (other than CLCC1) that could be fusogen candidates. A nuclear envelope fusogen could be encoded by an essential gene, and if so, it would be lost from the CRISPR library during passaging. Alternatively, CLCC1 itself could remodel membranes, effecting their fusion. CLCC1 does not resemble any known membrane fusogens ^40^. However, it has two conserved amphipathic helices that could, in principle, interact with membranes. One of them is a long bow-shaped AH that is positioned to interact peripherally with the membrane (**Fig 4a**). The other is an amphipathic helix FD_h2_ – the knuckle part of the fist domain (FD) – that is oriented perpendicular to the TMs (and parallel to AH) and juts outward (**Fig 4b and Extended Data Fig 7**). The membrane-distant surface of the helix FD_h2_ has several aromatic residues (W260, W265, F268, and W272), several of which are conserved (**Extended Data Fig 7)**. This is reminiscent of fusion peptides of class I viral fusogens – membrane-interacting spans that are typically enriched in aromatic and aliphatic residues ^40^. In its membrane-distant location, FD_h2_ is positioned to interact with the opposing membrane. Fusion peptides of some viral fusogens, e.g., Ebola virus GP, are stabilized by disulfides ^40^, just like helix FD_h2_. Future studies will clarify the fusion-promoting mechanism of CLCC1.

In addition to its role in membrane fusion, CLCC1 could facilitate nuclear egress in other ways by interacting with host or viral binding partners. One appealing idea is that the ER(PNS)-facing FD could function as a receptor for PEVs at the ONM. In this role, CLCC1 would act to promote fusion of PEVs with the ONM – and thus translocation of capsids into the cytoplasm – thereby preventing them from fusing with the INM, which would result in a counterproductive “back-fusion” releasing capsids back into the nucleus.

While the precise mechanism by which CLCC1 promotes fusion of the nuclear envelope remains undiscovered, collectively, our findings illuminate an ancient cellular membrane fusion mechanism important for nuclear envelope morphogenesis that herpesviruses co-opt for capsid nuclear egress.

## METHODS

### Antibodies

Mouse monoclonal antibody 8F5 ^17^ was produced by Cell Essentials, Inc. from a hybridoma generated by Dr. Jay Brown (University of Virginia) and provided by the University of Virginia Stem Cell Core Facility. Alexa-488-conjugated goat anti-mouse secondary antibody was purchased from Thermo Scientific (cat #A28175). Rabbit anti-CLCC1 polyclonal antibody was purchased from Sigma (cat #HPA009087). Rabbit anti-beta-actin monoclonal antibody was purchased from ABclonal (cat #AC026). Rabbit anti-Bip polyclonal antibody was purchased from Proteintech (cat #11587-1-AP). IRDye-800 conjugated goat anti-rabbit antibody was purchased from Li-Cor (cat #926-32211).

### Plasmids and cloning

BACmid of HSV-1-tdTomato with a UL34 deletion (BAC_GS3217-d34) was generated by En Passant Mutagenesis ^41^. The entire UL34 coding sequence was replaced by a start and stop codon. Sleeping beauty system plasmid and transposase plasmid ^42^ were purchased from Addgene (pSBbi-Hyg, Addgene 60524; pCMV (CAT)T7-SB100, Addgene 34879). MLF2-GFP plasmid ^28^ was a gift from Dr. Christian Schlieker (Yale University).

### Cell culture and maintenance

HeLa cells (ATCC CCL-2), Vero cells (ATCC CCL-81), PK15 (ATCC CCL-33), HEK293T (ATCC CRL-3216) were grown in Dulbecco’s modified Eagle medium (DMEM, Lonza) supplemented with 2 mM L-glutamine (Corning), 10% heat-inactivated fetal bovine serum (HI-FBS; Gibco), and 1X penicillin-streptomycin solution (Corning) at 37 °C, 5% CO_2_. Vero UL34 complementing cells (tUL34CX) ^43^ (a gift from Rich Roller, University of Iowa) were grown in the same medium but supplemented with 400 μg/mL G418 (Selleck Chemicals) every other passage. UL34-complementing cells containing Cre recombinase (Cre_tUL34CX) were generated by infecting Adenovirus (Ad5CMVCre-eGFP, University of Iowa) into tUL34CX.

### Virus strain and propagation

HSV-1 strain GS3217 is a strain F derivative that encodes an NLS-tdTomato transgene under the control of a CMV immediate-early (IE) promoter in place of the envelope glycoprotein gJ ^18^. PRV strain GS7741 is a strain pBecker3 derivative that encodes mCherry-NLS transgene under control of an MCMV IE promoter in place of the US2 gene. These two strains and HSV-2 strain 186 were gifts from Greg Smith (Northwestern University). For virus propagation, Vero or PK15 cells were seeded in a T175 flask at 1×10^7^ cells per flask on day 1. On day 2, cells were infected at MOI 0.01, and supernatant was harvested once the cytopathic effect (CPE) reached 100%, 72 hours post infection in general. Next, virus was pelleted by centrifugation at 41,000 g for 40 min at 4 °C, resuspended in the Opti-MEM medium (Gibco, cat #31985088) containing 10% glycerol (Chem-Impex, cat #30144), and stored at -80 °C for future use.

HSV-1 F strain GS3217-d34 containing UL34 deletion was made by transfecting the BAC_GS3217-d34 into Cre_tUL34CX. After the transfection, the cells were covered with 0.75% methylcellulose (Sigma) containing medium, then supernatant was collected from a single plaque forming area and subsequently propagated in the tUL34CX cells.

### Virus titration

For HSV-1 strain GS3217 and HSV-2, plaque assays were performed with HeLa and Vero cells. For PRV strain GS7741, plaque assays were performed in HeLa and PK15 cells. Briefly, HeLa or Vero or PK15 cells were seeded into 12-well plates on day 1 at 200,000 cells/well, on day 2, stock virus or supernatant was diluted in a series of dilutions and incubated with either Hela or Vero or PK15 cells for 1 h. Media was replaced by 0.75 methylcellulose (Sigma) containing DMEM medium. 3 days post infection, medium was aspirated out, cells were fixed and stained by 1% crystal violet (Sigma) in 50%/50% of methanol (Fisher Scientific)/water solution. Plaque forming unit per mL (PFU/mL) is quantified and calculated. For HSV-1 F strain GS3217-d34, HeLa cells were seeded in 96-well plate on day 1 at 15,000 cells/well. On day 2, the virus was serially diluted and added to cells. On day 3, tdTomato+ cells were counted using fluorescence microscope, and the titer was calculated as infectious units per mL (IU/mL).

### Multiple-step viral growth curves

Two control cell lines (HeLa and Int_4), two CLCC1-KO cell lines (cko3_4 and cko6_1), and two CLCC1-R cell lines (cko3_4_R_1 and cko6_1_R_1) were infected with HSV-1 (GS3217), HSV-2 strain 186, or PRV (GS7741) at MOI of 0.1. Supernatants were collected 24, 48, 72 hours post infection and frozen at -80 °C. Subsequently, plaque assays were performed in Vero cells to titer all the supernatants.

### Flow-cytometry-based nuclear egress assay

HeLa, intergenic region targeting, CLCC1 knockout, CLCC1 rescue or CLCC1 mutant rescue cell lines were seeded in 6-well plates on day 1 at 400,000/well. On day 2, cells were infected by the desired virus (HSV-1 strain GS3217 at MOI of 5, HSV-1 strain GS3217-d34 at MOI of 10). 24 h post infection, cells were trypsinized with 0.05% trypsin (Cytiva, SH30236.01) and collected by centrifuging at 500 g for 5 min. Cells were then fixed with 4% paraformaldehyde (PFA) (Thermo Scientific, J19943K2) for 1 h at room temperature before permeabilization with either 40 μg/mL digitonin or 0.2% TritonX-100 dissolved in PBS (Invitrogen, BN2006) for 20 min at room temperature. Next, cells were blocked by 0.5% BSA (Fisher Scientific BP1600100) for 1 h at room temperature, then incubated with capsid-specific 8F5 primary antibody (1:2000) for 1 h at room temperature or overnight at 4 °C. The cells were then washed with PBS and incubated with an Alexa488-conjugated secondary antibody (1:500) for 1 h at room temperature. Nuclear egress was measured by flow cytometry and quantified as the double-positive population (tdTomato+ indicating infection and Alexa488+ indicating capsids in the cytosol) in the digitonin permeabilized samples relative to the double positive population in the Triton X-100-permeabilized samples. Gating was based on HeLa-Cas9 cells infected with HSV-1 strain GS3217-d34 mutant strain, which has no nuclear egress, resulting in most of the population being tdTomato+/Alexa488-. Results were normalized to the nuclear egress of desired control cells (typically HeLa or Int_4) to calculate normalized nuclear egress percent.

### Generation of HeLa-Cas9 cell line

Lentiviral vectors pXPR_111 (Cas9), pXPR_047 (GFP and sgGFP), and pRosettav2 (antibiotic control) were provided by the Genetic Perturbation Platform group at the Broad Institute. HeLa-Cas9 cells were generated by infecting HeLa cells with pXPR_111, selecting with 10 mg/mL blasticidin (A.G. Scientific) for 2 weeks, and then maintaining with the treatment of blasticidin. Cas9 activity was tested by infecting the HeLa-Cas9 cells with pXPR_047. Cells were then selected with 2 mg/mL puromycin for 1 week, and % GFP+ cells were counted by flow cytometry. The HeLa-Cas9 cells with less than 25% GFP signal were used for generating Gattinara library cells.

### Generation of the Gattinara library HeLa cells

Lentiviral sgRNA Gattinara library (CP0073) targeting the whole genome (19993 target genes, 40964 sgRNAs) was provided by the Genetic Perturbation Platform group at the Broad Institute. The viral titer of the library is ∼1×10^8^ viral particles (VPs)/mL. Gattinara library HeLa cells were generated by infecting 1.5×10^8^ HeLa-Cas9 cells with Gattinara library lentivirus (Broad Institute, CP0073) with the amount of virus that allows 30% of cells to survive selection with 2 μg/mL puromycin (A.G. Scientific) for 1 week. After the selection, cells were maintained in the presence of puromycin-containing medium. Gattinara library lentivirus was titrated on HeLa-Cas9 cells. Briefly, lentivirus was first serially diluted. Next, 1 mL of 1.5×10^6^ HeLa-Cas9 cells and 1 mL of lentivirus was mixed in the present of 4 μg/mL polybrene, subsequently, the 2-mL mixture was put in one well of the 12-well plate, and spun down at 900 g for 1.5 h. The next day, 2 μg/mL of puromycin was used for selection. 7 days post selection, cells were collected, and cell viability assay was performed. The amount of lentivirus with 30% survival rate compared with non-infected group was used for library cell generation.

### CRISPR screen

1.5-2.5×10^8^ Gattinara library HeLa cells or 1×10^7^ HeLa cells were seeded in 10 cm dishes at 1×10^7^ cells/dish. On day 2, Gattinara library HeLa cells were infected with HSV-1 F strain GS3217 at an MOI of 5. As a control, HeLa cells were infected with HSV-1 F strain GS3217-d34 at an MOI of 10. Following infection, the cells were treated according to the flow-cytometry-based nuclear egress assay procedure outlined below. During sorting, ∼5-10% of tdTomato+/Alexa-488-Gattinara library HeLa cells were collected as cells without nuclear egress, and ∼70-85% of tdTomato+/Alexa-488+ Gattinara library HeLa cells were collected as cells with nuclear egress. DNA was isolated from both groups using the Qiagen Blood DNA kit per manufacturer’s protocol, except that fixed cells were incubated with proteinase K at 65 °C overnight instead of at 70 °C for 10 mins. The extracted DNA was sent to the Broad Institute for sequencing. Two independent Gattinara library transductions of HeLa-Cas9 cells were done (2 biological replicates), each with three independent HSV-1 infections (3 technical replicates), for a total of 6 experiments.

### Generation of CLCC1 knockout cell lines

Lentiviral vectors encoding sgRNAs targeting genes of interest or intergenic regions as controls were generated using a 2^nd^ generation, three-plasmid system consisting of pRDA_118 (sgRNA-containing plasmid), psPAX2 (packaging plasmid), and pMD2.G (VSV G envelope protein plasmid). pRDA_118 and pMD2.G were provided by the Genetic Perturbation Platform group at the Broad Institute. psPAX2 was a gift from Dr. Alexei Degterev (Tufts University). sgRNAs used to knockout CLCC1 were sgRNA-3: AGCTGTGGACATATGTACGT and sgRNA-6: TGTGTGCCAAAAAGATGGAC. The control intergenic site targeting sequence was ACAAAGGACCCCGGCGAAAG. All sgRNAs were inserted by Gibson assembly into the pRDA-118 backbone to make the plasmids (pRDA118_sgCLCC1_3), (pRDA118_sgCLCC1_6) and (pRDA118_sgInt).

HEK293T were transfected with the three plasmids using Genjet transfection reagent. After 24 h and 48 h, the lentivirus-containing supernatant was collected and stored either at 4 degrees C (short term) or -80 degrees C (long term).

Bulk CLCC1 knockout cells (cko3_bulk, cko6_bulk) and bulk intergenic site targeting cells (Int_bulk) were generated by infecting HeLa-Cas9 cells with lentivirus targeting CLCC1 (Lenti_sgCLCC1_3 or Lenti_sgCLCC1_6) or an intergenic-site (Lenti_sgInt), followed by selection with 2 μg/mL puromycin for one-week, and maintained in the puromycin containing medium. Single cell clones (cko3_2, cko3_4, cko6_1, cko6_2, Int_3, Int_4) were made by collecting single cells from the bulk population by single cell sorting with a flow cytometer, then expanding.

### Generation of CLCC1 rescue cell lines

IEF1a-CLCC1-CR plasmid was generated by inserting the CLCC1 gene synthesized by GenScript into pSBbi-Hyg with following changes to the sgRNA targeting sequences (CATGTGCTGAGACATATAGG to CACGTTCTTCGTCACATTGG, CATAGTTAAGCATGTCTGTG to CGTAATTCAGCATATCGGTC, AGCTGTGGACATATGTACGT to AACTTTGGACCTACGTGCGC, ATTATATGGATCCACTCCAA to GTTGTACGGGTCAACGCCGA, GCATATTGGAAAAGGAACTG to ACACATCGGCAAGGGCACCG, and TGTGTGCCAAAAAGATGGAC to TTTGCGCGAAGAAAATGGAT). hPGK1-CLCC1-CR plasmid was generated by switching the EIF1-alpha promoter to hPGK1 promoter (GeneScript). Bulk CLCC1 rescue cell lines (Int_4_IEF1a, Int_4_R_bulk, cko3_4_IEF1a, cko3_4_R_bulk, cko6_1_IEF1a, cko6_1_R_bulk) were generated by co-transfecting 1 μg of IEF1a-CR-CLCC1 or hPGK1-CR-CLCC1 with 100 ng of pCMV (CAT)T7-SB100 into cko3_4 or cko6_1 cells using GenJet transfection reagent (SignaGen). Cells were selected for two weeks with 300 μg/mL hygromycin. Single clones (cko3_4_R_1, cko3_4_R_6, cko6_1_R_1, cko6_1_R_6) were made by single cell sorting, and subsequently expanded.

All point mutations, D152R/D153R, E175R/D176R, D181R, D25E/D181R, S263R, W267R, D277R, K298E were generated in the hPGK1-CLCC1-CR backbone (GenScript). Bulk CLCC1 mutant rescue cell lines were generated by co-transfecting 1 μg of hPGK1-CR-CLCC1 mutants (D152R/D153R, E175R/D176R, D181R, D25E/D181R, S263R, W267R, D277R, K298E) with 100 ng of pCMV (CAT)T7-SB100 into Int_4 or cko6_1 cells using GenJet transfection reagent and selecting cells for 2 weeks with 300 μg/mL hygromycin. Cell lines were maintained in hygromycin-containing media.

### ER stress induction and cell viability assay

To induce ER stress, HeLa cells were treated with DTT at 1.5 mM for 4 h. Subsequently, cells were either non-infected or infected with HSV-1 (GS3217), and cells are maintained in the presence of 0.38 mM of DTT to maintain the stress level. 24 h post infection, Bip levels were measured by western blot. The cell viability was tested by treating HeLa or cko6_1 cells with different amounts of DTT for 24 hours, and measuring with Cell-titer Glo 2.0 (Promega), according to manufacturer’s protocol.

### Western Blot analysis

Cells were washed with cold PBS, lysed with RIPA buffer, and the lysates were spun at 14,000 g. Then, the supernatants were collected, and the total protein concentration was measured by BCA assay and normalized across samples for each experiment. Next, samples were mixed with SDS sample buffer, incubated at 95 °C for 5 minutes, and loaded into SDS-PAGE gel (Bio-Rad, cat# 456-1086). The proteins were transferred onto a nitrocellulose membrane (GE Healthcare, cat # 10600002) using the Trans-Blot Turbo Transfer System (Bio-Rad). The blot was then blocked with 5% milk in TBST buffer for 1 hour at room temperature, incubated with primary antibody (CLCC1 1:2000; Bip 1:1000; actin 1:1,000,000) overnight at 4 °C, washed with TBST 3 times, and incubated with goat anti-rabbit secondary antibody (1:5000) for 1 hour at room temperature. Images were collected on a LI-COR imager.

### Confocal Microscopy

Two control cell lines (HeLa and Int_4), two CLCC1-KO cell lines (cko3_4 and cko6_1), and two CLCC1-R cell lines (cko3_4_R_1 and cko6_1_R_1) were seeded at 75,000 cells/well in a 24-well plate (Greiner Bio-One, 662160) with a glass coverslip in each well (Chemglass, CLS-1760-012). The next day, cells were infected by either HSV-1 GS3217 at MOI of 5 or HSV-1 GS3217-d34 at MOI of 10. At 24 h post infection, cells were fixed by 4% PFA at room temperature for 20 minutes and either partially permeabilized by incubation in 40 μg/mL digitonin in PBS or fully permeabilized by 0.2% Triton X-100 for 20 minutes at room temperature. Cells were subsequently blocked with 0.5% BSA for 1 hour, incubated with 1: 2000 8F5 primary antibody overnight at 4 °C, washed 3 times with PBS, incubated with Alexa-488-conjugated goat anti-mouse secondary antibody diluted 1:500 for 1 hour at room temperature, washed 3 times with PBS, and stained with DAPI diluted 1:1000 for 5 minutes at room temperature. Cells were imaged using Leica SP8 confocal microscope.

### Transmission Electron Microscopy

HeLa, Int-4, cko3-4, cko6-1, cko3_4_R_1 and cko6_1_R_1 cell lines were either mock infected or infected with HSV-1 F strain GS3217. 24 h post infection, cells were collected, fixed, stained with osmium tetroxide and uranyl acetate, then embedded into resin. Subsequently samples were cut into thin slices and stained with lead citrate. EM images were collected on Morgagni or Tecnai electron microscopes.

### Multiple sequence alignments

The sequences of CLCC1 homologues from different species were obtained from the NCBI GenBank: Homo sapiens NP_001041675.1 (Human_CLCC1), Mus musculus NP_001171242.1 (MOUSE_CLCC1), Danio rerio XP_009294671.1 (DANRE_CLCC1), Xenopus tropicalis NP_001081605.1 (XENLA_CLCC1), OsHV-1 ASK05584.1 (OsHV1_ORF57), AbHV-1 AET44204.1) (AbHV1_ORF90), IctHV1 QAB08501.1 (IctHV1_ORF16a), AngHV QRM16927.1 (AngHV1_ORF112), Crassostrea gigas (Pacific oyster) XP_011439362.3 (Oyster_CLCC1), Haliotis rubra (blacklip abalone) XP_046552278.1 (Abalone_CLCC1), Ictalurus punctatus (Channel catfish) XP_047014136.1 (Ictalurus_CLCC1), Anguilla rostrata (American eel) XP_064188152 (Anguilla_CLCC1).

### Protein structure and topology predictions

Structural models of full-length, monomeric human CLCC1, OsHV-1 ORF57 and IcHV1 ORF16a, as well as hexameric, decameric, or hexadecameric human CLCC1 (residues 161 to 361) and OsHV-1 ORF57 (residues 51 to 250) were generated using the AlphaFold 3.0 online server (https://golgi.sandbox.google.com/) ^30^. Transmembrane domains were predicted by TMHMM 2.0 (https://services.healthtech.dtu.dk/services/TMHMM-2.0/).

### Statistical analysis

All the statistical analyses were done in GraphPad Prism 10. For the volcano plot, significance was calculated using paired multiple t-test. For the flow-cytometry-based nuclear egress assay data, significance was calculated using one-way ANOVA, with multiple comparisons.

## Supporting information

Supplementary Information (Supplementary Figures 1-2, Supplementary Tables 1-3, Supplementary References)

## ACKNOWLEDGEMENTS

We thank Samantha Moores for help with the development of the flow cytometry nuclear egress assay. We thank Dr. Greg Smith (Northwestern University) for the gift of bacmids and viral strains, Dr. Rich Roller (University of Iowa) for the gift of UL34-complementing Vero cell line, Dr. Alexei Degterev (Tufts University) for the gift of psPAX2 plasmids, Dr. Jay Brown (University of Virginia) and the University of Virginia Stem Cell Core Facility for the 8F5 antibody and the hybridoma cell line, and Dr. Christian Schlieker (Yale University) for the gift of MLF2-GFP plasmid. We are grateful to the staff at the Broad institute for providing reagents and advice. We are grateful to Drs. Stephen Kwok and Allen Parmelee at the David Thorley-Lawson Memorial Flow Cytometry Core Facility for help with the flow cytometry. We thank Dr. Berith Isaac (Brandeis Electron Microscopy Facility) and Dr. Maria Erickson (Harvard Medical School Electron Microscopy Facility) for help with transmission electron microscopy experiments. We thank Drs. John Coffin, Karl Munger, Ralph Isberg (Tufts University), and Marta Gaglia (University of Wisconsin-Madison), for helpful discussions.

Flow cytometry was performed at the David Thorley-Lawson Memorial Flow Cytometry Core at Tufts University School of Medicine, which is supported by NIH grant S10OD032201 (Stephen Kwok). Confocal microscopy was performed at the Imaging and Cell Analysis Core Facility within the Center for Neuroscience Research at Tufts University School of Medicine, which is supported by NIH grant P30 NS047243 (Rob Jackson). TEM samples were prepared and imaged at the Brandeis Electron Microscopy Facility and the Harvard Medical School Electron Microscopy Facility.

Research reported in this publication was supported by the National Institutes of Health under Award Number R01AI147625 (E.E.H.), and by a Faculty Scholar grant 55108533 from the Howard Hughes Medical Institute (E.E.H.). The content is solely the responsibility of the authors and does not necessarily represent the official views of the National Institutes of Health.

## AUTHOR CONTRIBUTIONS

B.D. designed the experiments, generated new reagents, carried out all the experiments, analyzed the data, generated hypotheses and models, and wrote the initial draft of the manuscript.

L.P. generated CLCC1 knockout cell lines, conducted the CLCC1 knockout and rescue experiments, and tested the ER stress effect.

A. S. tested the phenotypes of the CLCC1 mutants.

H. D. assisted with the CRISPR screens.

T. H. generated the CLCC1 rescue cell lines.

C. D. and C. L. generated CLCC1 knockout cell lines.

J. G. D. helped design and troubleshoot the CRISPR screen, provided reagents, and did sequencing for CRISPR screen.

E.E.H. oversaw all aspects of the project, designed the experiments, analyzed the data, generated hypotheses and models, acquired funding, and wrote the initial draft of the manuscript.

All authors edited and finalized the manuscript.

---

**Extended Data Figure 1.**
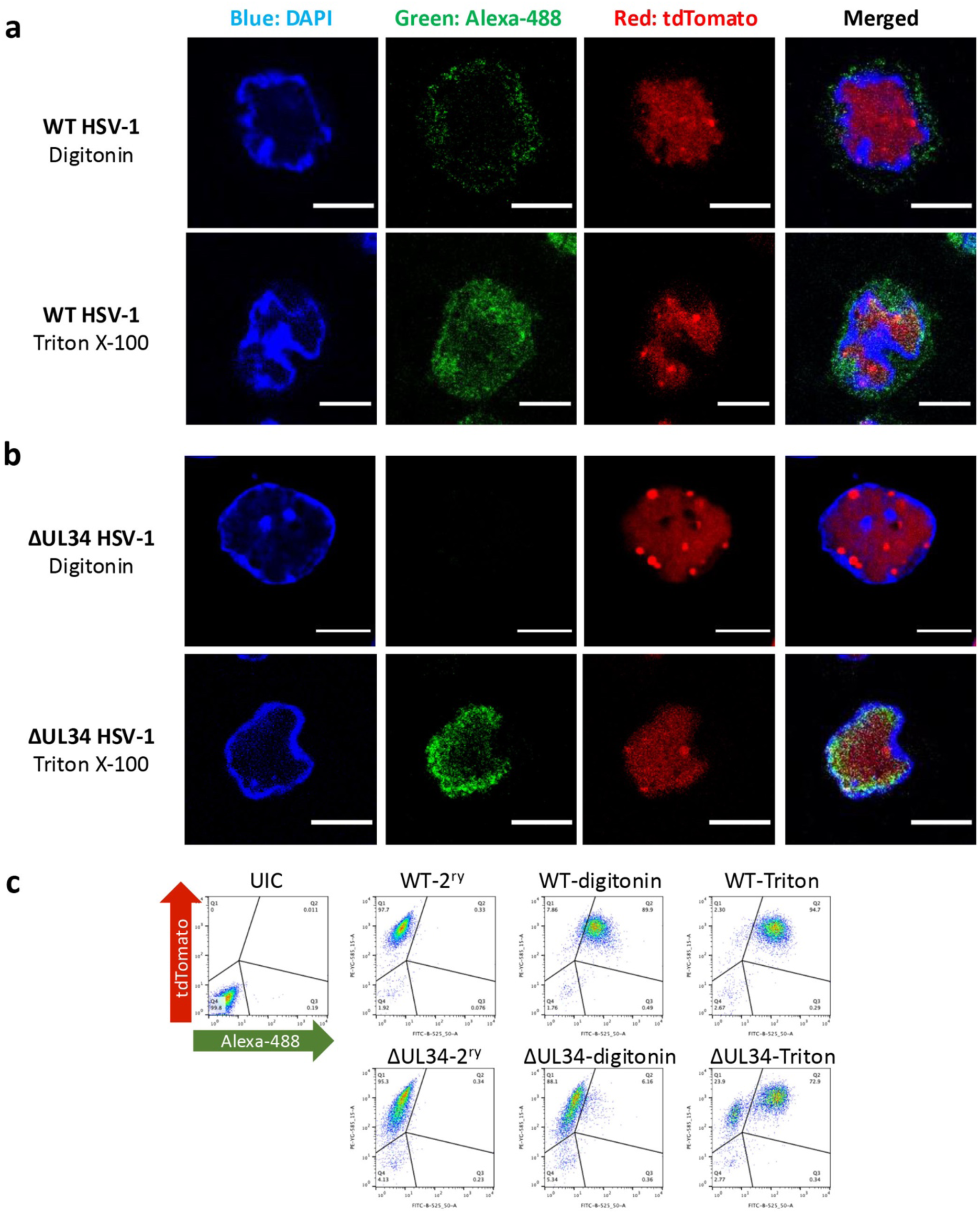
Development of the flow-cytometry based assay to measure nuclear egress. **a,b)** Confocal images of HeLa cells infected with either WT HSV-1 **(a)** or HSV-1 ΔUL34 mutant deficient in nuclear egress **(b).** Cells were either partially permeabilized with digitonin (top) or fully permeabilized with Triton X-100 (bottom) and then stained with a capsid-specific primary antibody and Alexa-488-conjugated secondary antibody (green). Nucleus was stained with DAPI (blue). tdTomato signal indicates infection (red). Scale bar = 10 mm. **c)** Flow cytometry data for uninfected HeLa cells (UIC), or HeLa cells infected with WT HSV-1 (top) or HSV-1 ΔUL34 mutant. Infected cells were either partially permeabilized with digitonin or fully permeabilized with Triton X-100 and the stained with capsid-specific primary antibody and Alexa-488-conjugated secondary antibody. “2^ry^” samples were partially permeabilized with digitonin and incubated with secondary antibody only. Each image is a representative of three biological replicates.

**Extended Data Figure 2.**
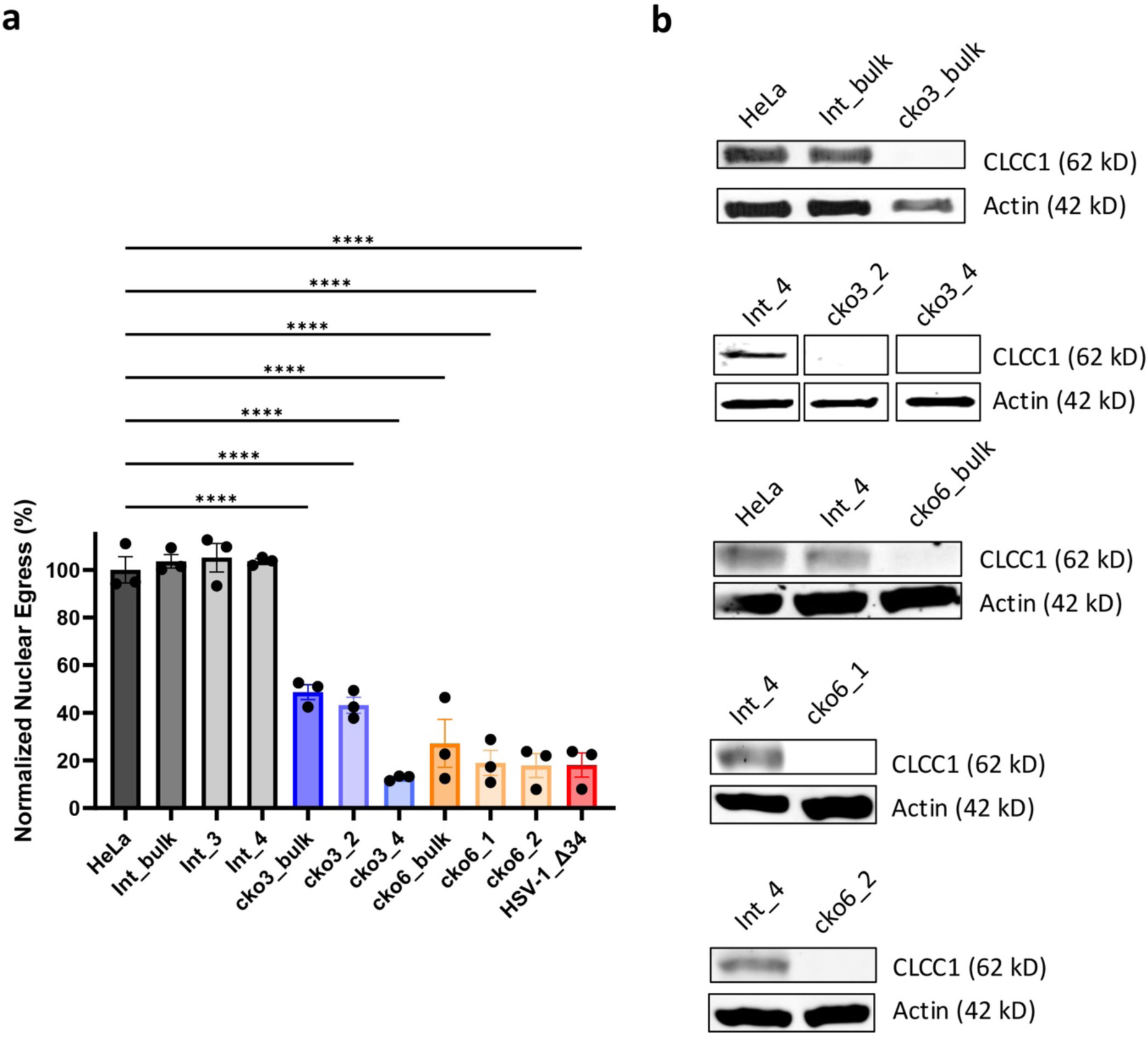
CLCC1 depletion causes a defect in HSV-1 nuclear egress. **a)** Bulk CLCC1-KO (cko3_bulk and cko6_bulk), single-clone CLCC1-KO (cko3_2, cko3_4 and cko6_1, and cko6_2), HeLa, bulk intergenic site targeting (Int_bulk), or single-cell intergenic site targeting (Int_3 and Int_4) cell lines were infected with WT HSV-1 at an MOI of 5. As a positive control, HeLa cells were infected with HSV-1 ΔUL34 mutant virus, defective in nuclear egress, at an MOI of 10. Nuclear egress was measured at 24 hpi by the flow cytometry nuclear egress assay and normalized to HSV-1-infected HeLa cells. Each experiment had three biological replicates, each containing two technical replicates. Each data point represents a biological replicate. Bars represent mean values, and the error bars represent SEM. P < 0.0001 = ****. Significance was calculated using one-way ANOVA, with multiple comparisons. Data for HeLa, Int_4, cko3_4, cko6_1, and HSV-1 ΔUL34 are the same as in **Fig 2a**. **b)** Western Blot analysis of CLCC1 levels in cell lines used in **(a).** Each image is a representative of three biological replicates. The bands in the second blot from the top were cut from the same gel and rearranged for better visualization.

**Extended Data Figure 3.**
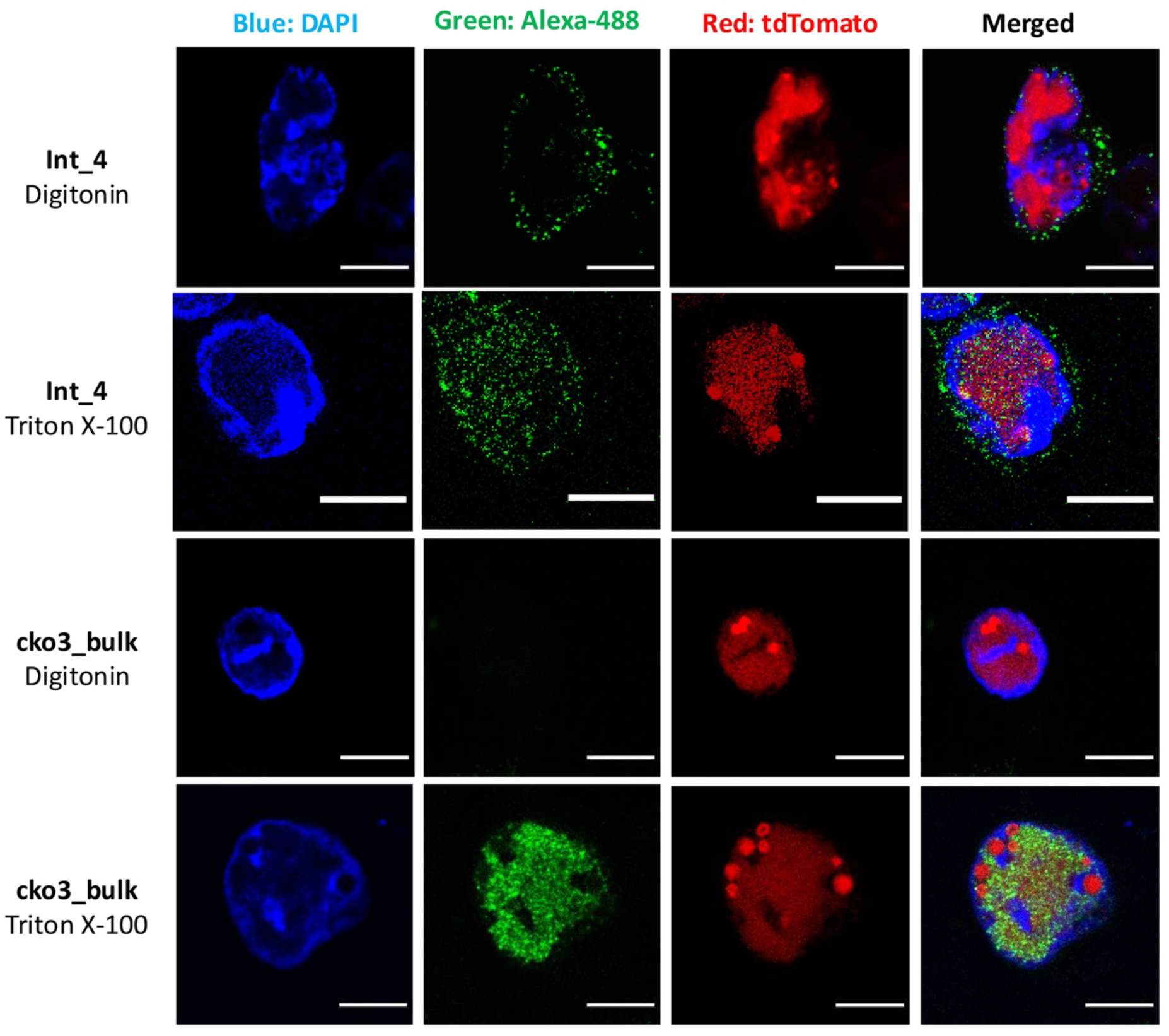
CLCC1 depletion causes a defect in HSV-1 nuclear egress, visualized by confocal microscopy. Confocal images of Int_4 and bulk CLCC1-KO (cko3_bulk) cell lines infected with WT HSV-1. Cells were either partially permeabilized with digitonin or fully permeabilized with Triton X-100 and then stained with a capsid-specific primary antibody and Alexa488-conjugated secondary antibody (green). Nucleus was stained with DAPI (blue). tdTomato signal indicates infection (red). Scale bar = 10 mm. Each image is a representative of one biological replicate.

**Extended Data Figure 4.**
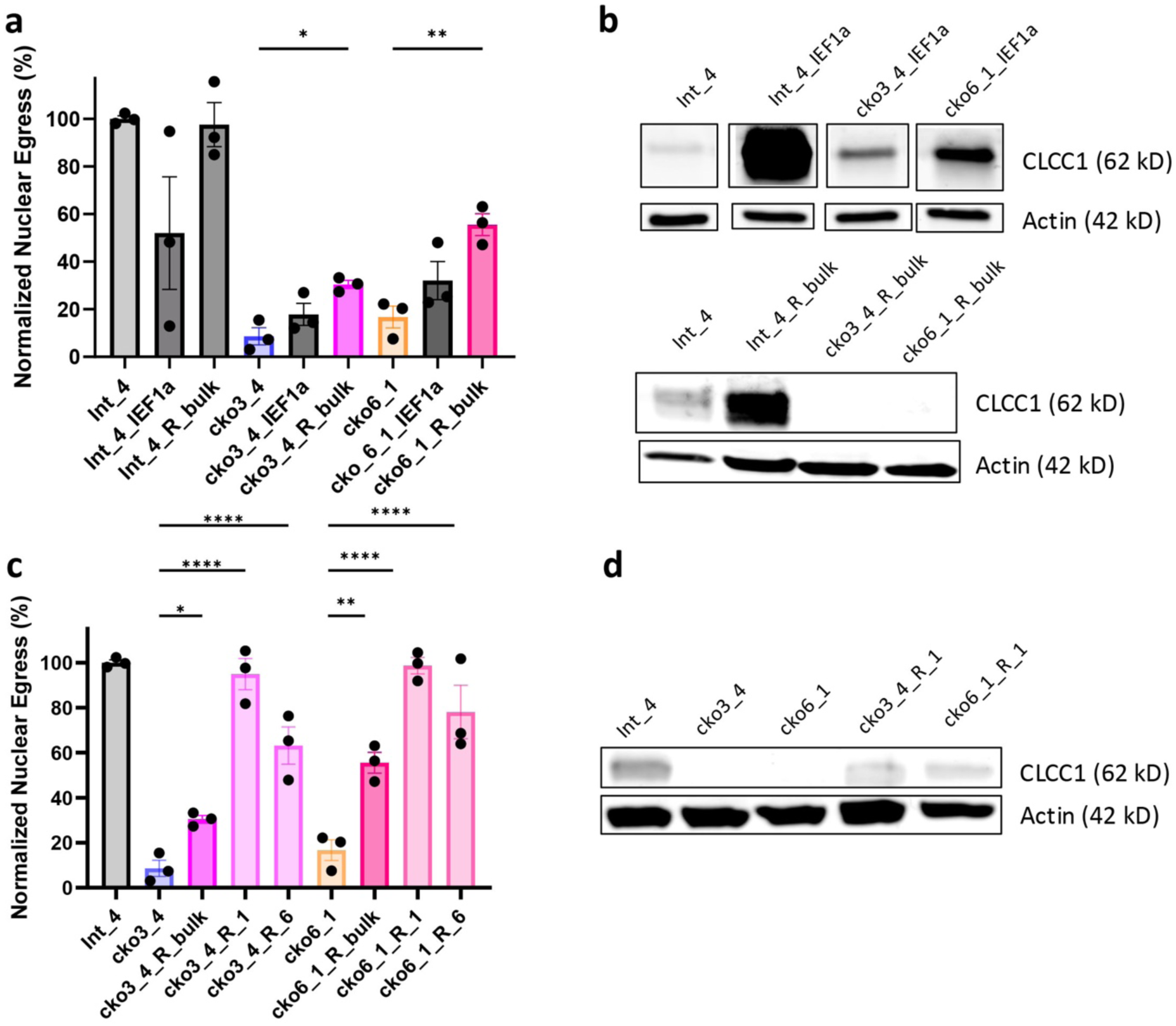
Expression of CLCC1 *in trans* can rescue the HSV-1 nuclear egress defect due to CLCC1 depletion. **a)** Single-clone CLCC1-KO (cko3_4 and cko6_1), CLCC1 overexpressing (Int_4_IEF1a and Int_4_R_bulk, with CLCC1 under the control of a strong and weak promoter, respectively), and bulk CLCC1-R (cko3_4_IEF1a, cko6_1_IEF1a, cko3_4_R_bulk, and cko6_1_R_bulk, with CLCC1 under the control of a strong weak promoter, respectively), HeLa, and single-cell intergenic site targeting (Int_4) cell lines were infected with WT HSV-1 at an MOI of 5. Nuclear egress was measured at 24 hpi by the flow cytometry nuclear egress assay normalized to HSV-1-infected Int_4 cells. Each experiment had three biological replicates. Each data point represents a biological replicate. Bars represent mean values, and the error bars represent SEM. P < 0.0001 = ****, P < 0.01 = **, P < 0.05 = *. Significance was calculated using one-way ANOVA, with multiple comparisons. Data for HeLa, Int_4, cko3_4, and cko6_1 are the same as in **Fig 2a**. **b)** Western Blot analysis of CLCC1 levels in single-clone CLCC1-KO (cko3_4 and cko6_1), control Int_4, CLCC1 overexpressing (Int_4_IEF1a and Int_4_R_bulk), and bulk CLCC1-R (cko3_4_IEF1a, cko6_1_IEF1a, cko3_4_R_bulk, and cko6_1_R_bulk) cell lines. Each image is a representative of at least two biological replicates. The bands in the top blot were cut from the same gel and rearranged for better visualization. **c)** Single-clone CLCC1-KO (cko3_4 and cko6_1), bulk CLCC1-R (cko3_4_R_bulk and cko6_1_R_bulk), Intergenic-site targeting (Int_4), single-clone CLCC1-R (cko3_4_R_1, cko3_4_R_6, cko6_1_R_1, and cko6_1_R_6), and control Int_4 cell lines were infected with WT HSV-1 at an MOI of 5. Nuclear egress was measured at 24 hpi by the flow cytometry nuclear egress assay normalized to HSV-1-infected Int_4 cells. Each experiment had three biological replicates. Each data point represents a biological replicate. Bars represent mean values, and the error bars represent SEM. P < 0.0001 = ****, P < 0.01 = **, P < 0.05 = *. Significance was calculated using one-way ANOVA, with multiple comparisons. Data for HeLa, Int_4, cko3_4, and cko6_1 are the same as in **Fig 2a**. Data for Int_4, cko3_4, and cko6_1 are the same as in **Fig 2a**. Data for cko3_4_R_bulk and cko6_1_R_bulk are the same as in **(a). d)** Western Blot analysis of CLCC1 levels in single-clone CLCC1-KO (cko3_4 and cko6_1), single-clone CLCC1-R (cko3_4_R_1 and cko6_1_R_1), and control Int_4 cell lines. Each image is a representative of three biological replicates.

**Extended Data Figure 5.**
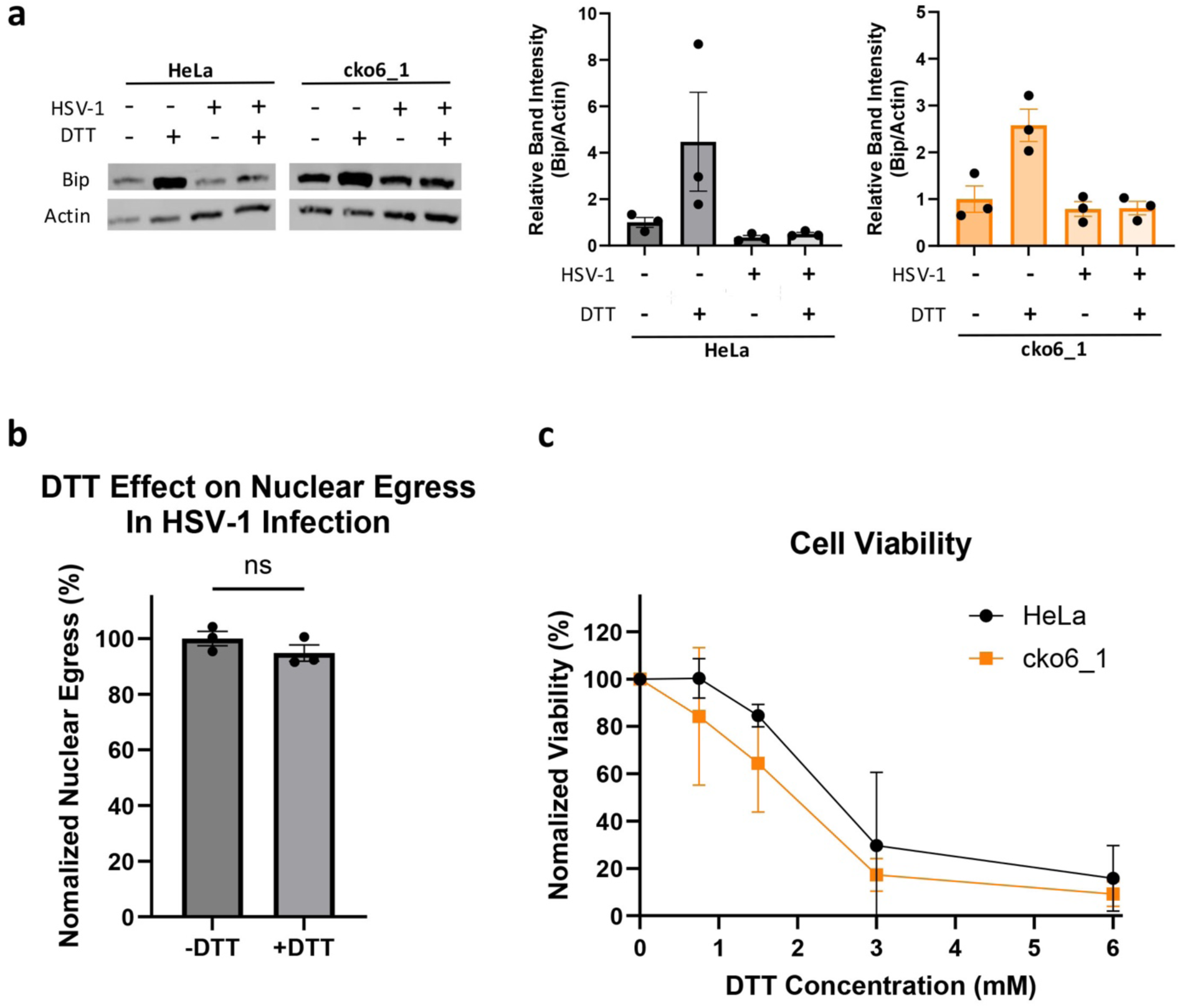
CLCC1 role in nuclear egress is independent of its role in ER stress response. **a)** Western Blot analysis of BiP (ER stress marker). *(Left)* HeLa or cko6_1 cells were either pre-treated with 1.5 mM DTT for 4 h or left untreated, and then either infected with WT HSV-1 at an MOI of 5 or uninfected. Following infection, cells pre-treated with DTT were incubated in the presence of 0.38 mM DTT for an additional 24 h. Each image is a representative of three biological replicates. *(Right)* quantifications of three replicate Western Blots, normalized to the HeLa or cko6_1 untreated and uninfected group. Each data point represents a biological replicate. Bars represent mean values, and the error bars represent SEM. **b)** HeLa or cko6_1 cells, treated as in **(a)**. Nuclear egress was measured at 24 hpi by the flow cytometry nuclear egress assay. Each experiment had three biological replicates, each with three technical replicates. Each data point represents a biological replicate. Bars represent mean values, and the error bars represent SEM. **c)** Cell viability of HeLa or cko6_1 cells, following treatment with different DTT for 24 hours. Viability was measured and normalized to HeLa or cko6_1 mock treatment group. Each symbol is the mean of three biological replicates, and bars show SEM.

**Extended Data Figure 6.**
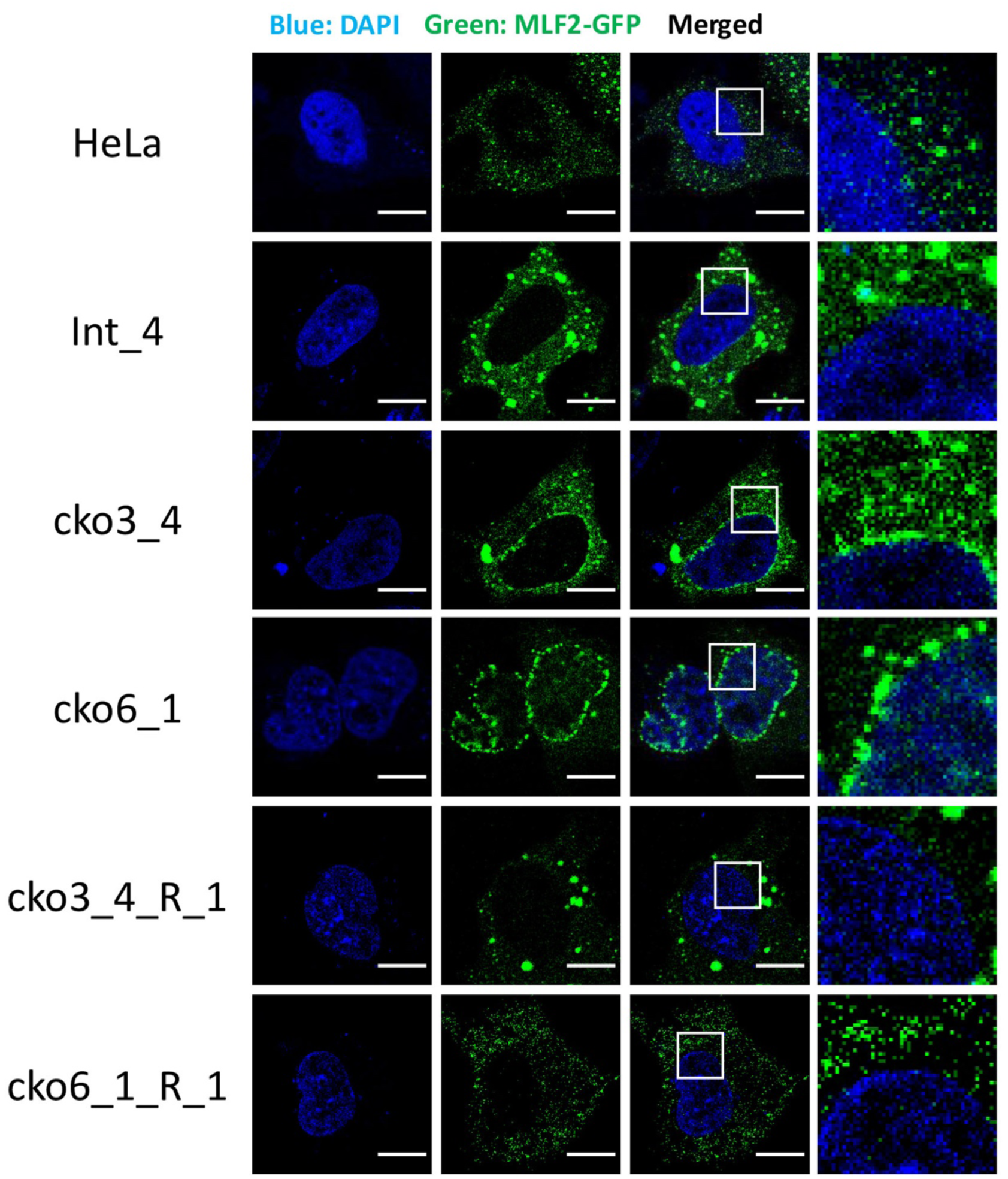
CLCC1 depletion causes MLF2 accumulation in the nuclear envelope, visualized by confocal microscopy. Confocal images of single-cell CLCC1-KO (cko3_4 and cko6_1), single-cell CLCC1-R (cko3_4_R_1 and cko6_1_R_1), and control HeLa and Int_4 cell lines overexpressing MLF2-GFP (green). Nucleus was stained with DAPI (blue). Scale bar = 10 mm. Each image is a representative of one biological replicate.

**Extended Data Figure 7.**
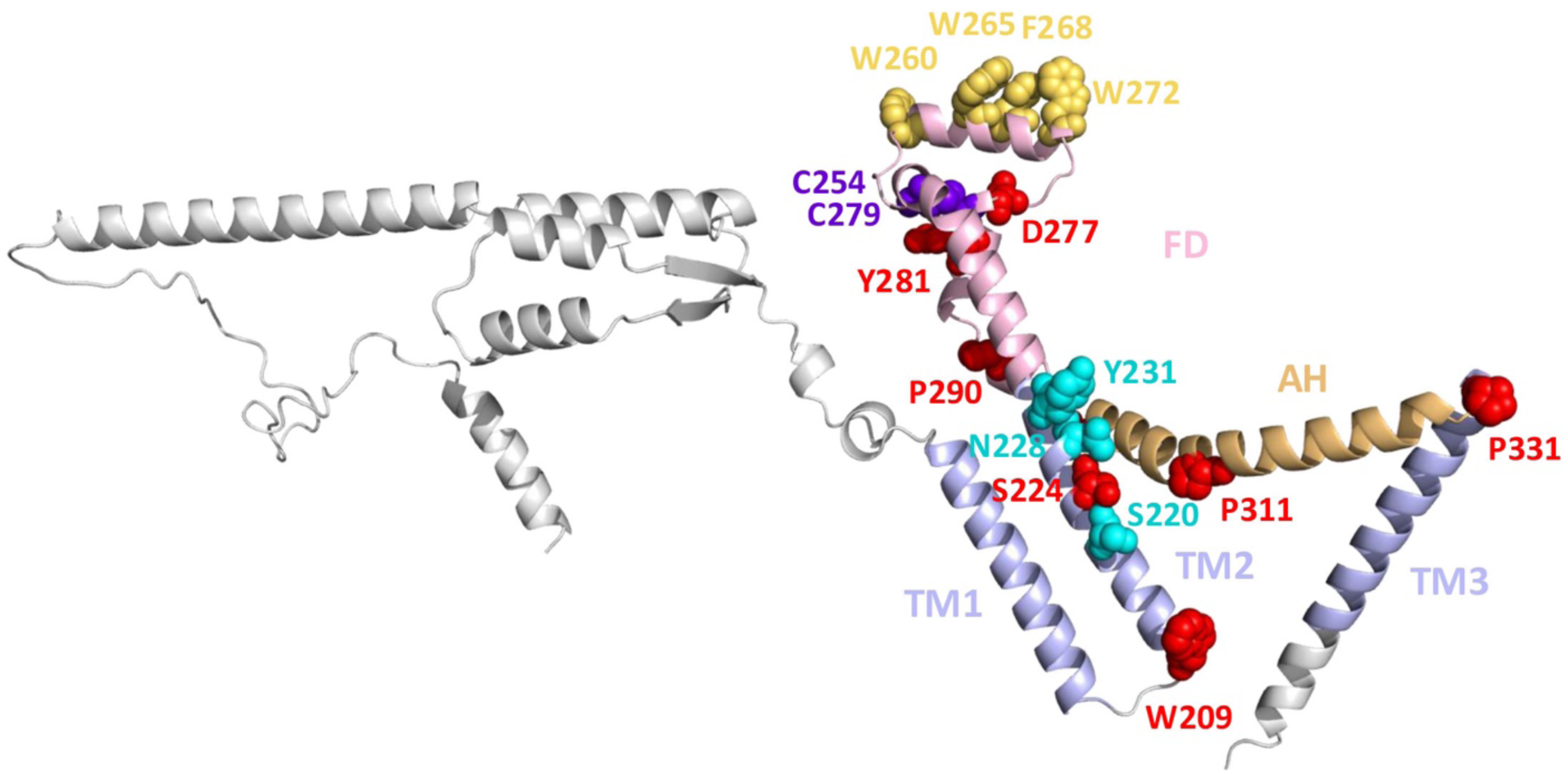
Residues of potential functional importance in human CLCC1. A ribbon diagram of an AlphaFold3 model of human CLCC1. Structural elements and domains are colored as in **Fig 5** and labelled. Mutated residues are shown in sphere representation and colored as follows: 10 residues that are invariant across 8 representative animal and 4 herpesviral homologs (red, except cysteines shown in purple), aromatic residues in the “knuckle” FD_h2_ helix of FD (yellow orange), polar spine residues in TM2 (cyan). Residues 365-539 were removed, for clarity.

**Extended Data Figure 8.**
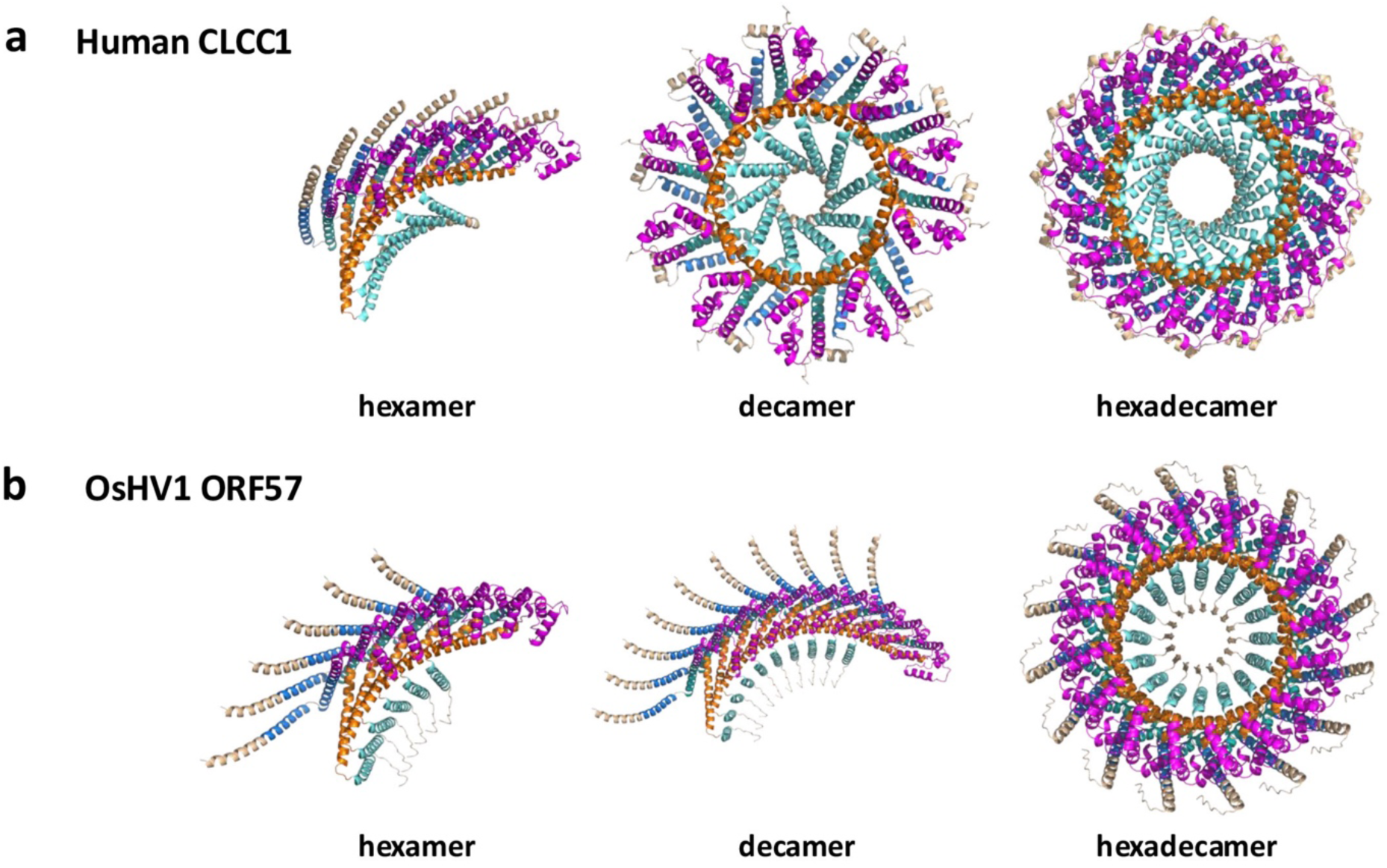
AlphaFold3 models of human and herpesviral CLCC1 multimers. **a)** AlphaFold3 models of the core region of human CLCC1 (residues 161-360) as a hexamer *(left)*, decamer *(middle)*, or hexadecamer *(right)*. b) AlphaFold3 models of the core region of OsHV-1 ORF57 (residues 66-268) as a hexamer *(left)*, decamer *(middle)*, or hexadecamer *(right)*. Structural elements and domains are colored as in **Fig 4**: TM1 (blue), TM2 (deep teal), FD (magenta), AH (orange), and TM3 (teal).

## Notes

### Competing Interest Statement

The authors have declared no competing interest.

