## Supplementary Information (Supplementary Figures 1-2, Supplementary Tables 1-3, Supplementary References) for "CLCC1 promotes membrane fusion during herpesvirus nuclear egress"

**The PDF file includes:**

Supplementary Figures 1-2
Supplementary Tables 1-3
Supplementary References

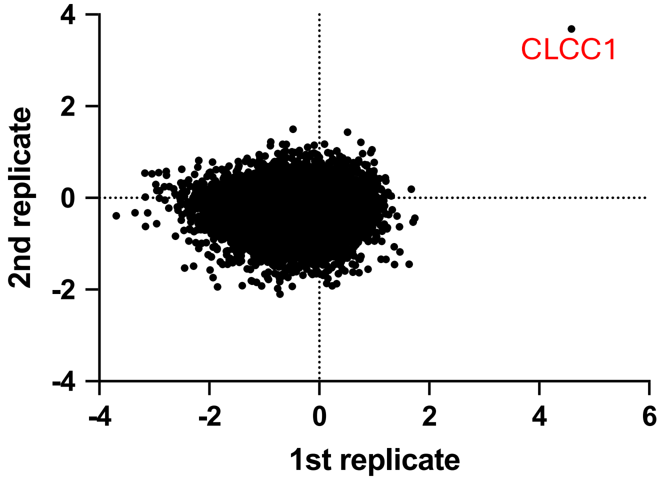


**Supplementary Figure 1. Genome-wide CRISPR screen results.** Comparison of the log fold changes between the 1st (x-axis) and 2nd (y-axis) biological replicate of the CRISPR screen. Each dot indicates one gene. The liner regression was calculated, yielding $R^2 = 0.0012$. Top hit, CLCC1, is labelled in red.

```
HUMAN_CLCC1       1  ...........MLCSLLLCECLLLV.....AGY.AHDDDWIDPTDMLNYDAASGTMRKSQ
MOUSE_CLCC1       1  ...........MLCRLLLCECLLLI.....TGY.AHDDDWIDPTDMLNYDAASGTMRKSQ
DANRE_CLCC1       1  ..MKLSSSSSFGLCILVVFFCFVVIESAKIRIDGYNDEAWIDPYDMLNYDPTTKRMRKST
XENLA_CLCC1       1  ............MRLFLLVALYLS.....PVYGDYTDEWIDPSDMLNYDAASGKMKNKP
OsHV1_ORF57          ............................................................
AbHV1_ORF90          ............................................................
IcHV1_ORF16A         ............................................................
AngHV1_ORF112        ............................................................
Oyster_CLCC1      1  ......MGSMRVWVFILLLWCAV........CVLMEEDVFIDPHDMVNPRRKILKEEG..
Abalone_CLCC1     1  .......MSITMWYMTVVLLCSVVQ..HCHTEVRMDTDDWIDPYDMVNFNHDTMTMKGKQ
Ictalurus_CLCC1   1  MKMKVSGGLLLRVCGGL......LVISSVLSFVQSEDEEWIDPYDMLNYDPASKSMRKPA
Anguilla_CLCC1    1  .........................................MLNYDASTKTMRKTE

HUMAN_CLCC1      44  AKYGI...................SGEKDVSPDLSCADEISECYHKLDSLT......Y
MOUSE_CLCC1      44  VRSGT...................SEKKEVSPDSSEAEELSDCLHRLDSLT......H
DANRE_CLCC1      59  ESESYQNVP.............TKRREFNSESCDVPKCPDEHECIKKLHILQ......K
XENLA_CLCC1      43  QVESTQSYYSVENTVSQDATQQPAQKANELHQNPDMTCSAEYQEYKTKLENLK......G
OsHV1_ORF57       1  ......................................................MTDAVK
AbHV1_ORF90       1  ......................................MSCKNHLEMLTRLAVRLT
IcHV1_ORF16A      1  ....................................MTGECFFITIIHGGHTKHMN
AngHV1_ORF112        ............................................................
Oyster_CLCC1     45  ..ETPEE....EKTLKI.NPKLPDIQPSISEDNTLLP...DIEDIKKPADVKQ.......
Abalone_CLCC1    52  EGGTPQA....NHNQDQGQRQVPKKDESITQETES.E....ASKMVEDAERSKG.......
Ictalurus_CLCC1  55  ESSSYSNVP.............TKRREYSQESCQVAQCPEVKECTDKVNMLQ......R
Anguilla_CLCC1   16  P..SHANGP.............TKRREYMLDSSQKPSCPDITEYSNKVASLQ......R

HUMAN_CLCC1      77  KIDECEKKKRE.DYESQSNPVFRRYLNKILIEAGKLGLPDE....NKGDMHY........
MOUSE_CLCC1      77  KVDSCEKKKMK.DYESQSNPVFRRYLNKILIEAGKLGLPDE....NKVEMRY........
DANRE_CLCC1      99  EFDEQKSKSTATLSKPVCLPVFKRFLSKLLKETSKLGLPDD....GITAMHY........
XENLA_CLCC1      97  QLE..ETKRME.KSKSKSQAIFKRYLNKILIEAGRIGLPDE....SYPKAHY........
OsHV1_ORF57       7  KIAKL............VVDLHGEKNTQNIIEVAKAGGKES..A...QAVLYKKLTDAII
AbHV1_ORF90      19  TLCKLKPDLECTTPA.SLLDLKSDFRKNPIS..AEEGEKSRPDEQNFVNNHFRLLFNSME
IcHV1_ORF16A     21  MMENVPALTMGDRTPCVYKTVLKRYLIRILKVVGDEGGEFE...................
AngHV1_ORF112        ............................................................
Oyster_CLCC1     88  .....DTPKECPNTD.IGKHLLRSYVKSLLGHF...ELKRP....FSGSQEY........
Abalone_CLCC1    97  .....DNPGTCSQPPSPCPALFRQYVKSLLFHM...KGK.V....ASGVEEL........
Ictalurus_CLCC1  95  EIEEIRRRETFSSQQPTCNPVFKRFLAKLLKEIDKLGLPTA....VTAEKHY........
Anguilla_CLCC1   54  EIQELEKRIASASQKPAIHPVFKRFLAKLMKLIKNLSLPKE....LHSDVHY........

HUMAN_CLCC1     124  .DAEIILKRETLLEIQKFLNGE.DW..KPGALDDALSDI...LINFKFHDFETWKWRFED
MOUSE_CLCC1     124  .DAEILLSRQTLLEIQKFLSGE.EW..KPGALDDALSDI...LINFKCHDSEAWKWQFED
DANRE_CLCC1     147  .DAEVKLSKQSLAEIQKLLNDEDGW..TTGAMDEALSQI...LVQFKLHDYEAWKWRFED
XENLA_CLCC1     142  .DAEVVFTMEMLQEIQSFLNNG.DW..NVGALDDALSST...LVQFKHHNEEEWKWKFED
OsHV1_ORF57      50  KDMTAMINHGDSKEKMQFTSTEQD...LPMAEDTTLTNN...............KLFWF..
AbHV1_ORF90      76  RDLKVIIESGATSSESPTASTDAE...PPR.LGQFVSND.............LPQTF..
IcHV1_ORF16A     62  .....VTSHNMERIKRLLGKWDES..SPGNL...EAV...LIDMKLTAAERPESDFSG
AngHV1_ORF112     1  ...MKMNRILVL.LLLFWGTRAEW..DVTSYKGLVKGC...KCRPEEVKEVKVWRFED
Oyster_CLCC1    127  .NMLLQLSVHDFEMLQKFVSEESKDLHTLHESAAILTDMIRSVSRSHLGTMGKISIWFED
Abalone_CLCC1   136  .NIRVKVSATEVEYLEQYTSSSSE..QNFHRVHEILSSMVQHVSEVSQDDPSVRRLWIEN
Ictalurus_CLCC1 143  .DAEVKLSKKRVIEIQKLVSDESSW..RTGALDDALSQI...LINFKHHDPETWKWRFED
Anguilla_CLCC1  102  .DAEVRLSRHAVAEIQKIIDEDDSL..RTGALDDALSKI...LVNFKQHDYEAWKWRFED

HUMAN_CLCC1     177  SFGVDPYNVLMVLLCLLCIVVLVATELWTYVRWYTQLRRVLIISFLFSLGWNWMYLYKLA
MOUSE_CLCC1     177  YFGVDPYNVFMVLLCLLCLVVLVATELWTYVRWYTQMKRIFIISFLLSLAWNWIYLYKMA
DANRE_CLCC1     201  TFHVDVDTVLKVSLIVLIIVAIICTQLWSVVSWFVQFRRMFAVSFFISLIWNWFHLYMLA
XENLA_CLCC1     195  SFGVDVYTLFMLILCVLCLVKLIATEIWTHIGWFTQLKRLLILSTVISFGWNWMYLYKVA
OsHV1_ORF57      91  ........SVVVG..IVLFVTAINYLLDKVCWIKAARKIAITCFLISVCWNYINIYEKT
AbHV1_ORF90     116  ........LLVAG..LLLVCVGLNYVLNRISWRFLIHRMCVVAFLASVIINFIHDYEKV
IcHV1_ORF16A    107  HFGVPRWEVILPILFVTMVVILIV....TTNRWTKWIPRLFACAFVMSLGWNWVYLLKMA
AngHV1_ORF112    51  TFGVEIDTVLQICSCVLVIVMIVCGELWSTVSWFLQLKRAFIICLFVSFIWNWIYLYKAA
Oyster_CLCC1    186  KYGASIDAALKMVAVLLMASVFATLEMKLQMSWRQRFMKLIVLSFIVSIPMTWFELYKAE
Abalone_CLCC1   193  VIGMQVEKFVMFALLAAALVTGMLVAARLHFSWKKLSMKLIFVLFVLSTVWQWVELYKIE
Ictalurus_CLCC1 197  TFGVEPDTVIKVSVIVLIIVVIICTEMWSLMSWFVQFKRMLAICFIISLVWNWFYLYKIA
Anguilla_CLCC1  156  TFGVEIGTVLQVCACVLIIVLIICGELWSSVSWFVQFRRVFAICFFVSIVWNWFYLYKIA

HUMAN_CLCC1     237  FAQHQAEVAKMEPLNNVCA.....KKMDWTGSIWEWFRS...SWTYKDDPCQKYYELLLV
MOUSE_CLCC1     237  FAQHQANIAGMEPFDNLCA.....KKMDWTGSLWEWFTS...SWTYKDDPCQKYYELLIV
DANRE_CLCC1     261  FAEHKKNIVQVESFNAKCTGL...KQLNWQDSLSEWYRR...TWTLQDDPCKKYYEVLVV
XENLA_CLCC1     255  FAERQAELAKLQDFD.KCS.....QKISWSESLFDWMKG...AATFQNDPCEDYFKALIV
OsHV1_ORF57     140  MAKRY..MVIRQGIPGGCMER....GADWSTVLRTMFKTLMITHSDSNDECLQYAKSIIV
AbHV1_ORF90     165  AAKRY..LKLRQGLPENCMER....GGSWRTVLSSLISYLFIITPEDQDECVTYAKAIVI
IcHV1_ORF16A    163  IAEHEAGLAKVDMAGLNCAR......DTWIGGLKEWFRT...TWTLEDDPCRKYHELLIV
AngHV1_ORF112   111  YAEHQANMIKLDGVAKRCAN......ADFMSTLKDWFRS...TWTLQDDPCKKYHEVLII
Oyster_CLCC1    246  QIKQE..TVAMKDAPAECIKNNDSISENWLNAGFQFITS...MVTLKEDKCQKYYEHVMI
Abalone_CLCC1   253  EAKQH..MAIMKEMPEGCRKRER..DEDFMSTVMRTLSS...TFTFEKDDCLKYYEHHLI
Ictalurus_CLCC1 257  FAEHQSNMVKMENVNGKCTGV...KKFDWMDNIKEWYRT...TLTLQEDPCKKYYEALIV
Anguilla_CLCC1  216  FAEHQTNIVKFESVNEKCTGV...KKIDWKDNLKEWFRS...TWTLQDDPCKKYYEVLMV
```

```
HUMAN_CLCC1      289  NPIWLVPPTKALAVTFTTFVTEPLKHIGKGTGEFIKALMKEIPALLHLPVLIIMALAILS
MOUSE_CLCC1      289  NPIWLVPPTKALAITFTNFVTEPLKHIGKGAGEFIKALMKEIPVLLQIPVLAILALAVLS
DANRE_CLCC1      315  NPILLVPPTKAITITITNFITDPLKHIGEGISEFLRALLKDLPVTLQIPVLIIIILAILI
XENLA_CLCC1      306  SPTLMVPPTKALALTFTNFITEPLKHIGKGIGEFLNALLSEIPLFFQVPVLIFIAVLLLA
OsHV1_ORF57      194  EPLFEVTPTTALATTISDLILVPINLAAKSCNNVFRTVFEGVPVFMIPILVFLLVYISTL
AbHV1_ORF90      219  EPLFEVTPMNAILTSVSTLVLTPVEQLAKSTNTIFKSLLYGIPSLLVPVVIGVILYVVTM
IcHV1_ORF16A     214  DPLLMVPPTRVLAMTCATFFMEPMRYVGSGIGLFIRELLGPLPVTLQLPVLIALVVFMAC
AngHV1_ORF112    162  NPVLLVPPTKAISVTFVTFVTEPLKHFGHGIGEFIKALLKDLPITLQVPVWGTILVAAAG
Oyster_CLCC1     301  DPFLKVPPTKAVGVTFVRFFLSPLKDVGASLSSFIRELLIDLPLTLYPVAMAMVTVFLFL
Abalone_CLCC1    306  APLWRATPTQALSRTFIRFFVSPLQDIGEALREFLVGLLKDLPAQLWPVALGGVALFFFL
Ictalurus_CLCC1  311  NPILLVSPMKAITMTIATFFTDPLKHIGQGISEFLRALLKDLPFTLQIPVLLLIGLSVVV
Anguilla_CLCC1   270  NPILLVPPTKAISVTITTFITEPLKHLGQGISEFLRALLKDLPVTLQIPVLLTIVLSILV

HUMAN_CLCC1      349  FCYGAGKSV....HVL..RHIGGPESEPPQ.ALRP....RDRRRQEEIDY.RPDGGA...
MOUSE_CLCC1      349  FCYGAGRSV....PML..RHFGGPDREPPR.ALEP....DDRRRQKGLDY.RLHGGA...
DANRE_CLCC1      375  FVYGSAQAAIHQVARF..PRLGWRQEQPPP.AVGQRQNPQLRAHEEP......WEGG...
XENLA_CLCC1      366  FFYGAGTAVMNPVNLY..RRLTGPEREKPL.PVEP....TRSNRKRFIEDVRVPPAL...
OsHV1_ORF57      254  CIISTKRYTISVPFLLE.LKPCL.........VEPPPMVPISVRNETCNVVHKKPVALRR
AbHV1_ORF90      279  IMICYNRYSVNFLSLLN.INPTAA.......LVHPAPPPPVVVTEKYVRARPKKRV...L
IcHV1_ORF16A     274  FGLALGKYSNRDIRTI..REIPVSHDGVPR.I.....................DGGVHRT
AngHV1_ORF112    222  FMYGISSAAAGHLSHFAQPHTTHTT...................HEDRRWKHFWKCGLDF
Oyster_CLCC1     361  ILFMTFGYSLRLPFFLS.IEPSPYHAVIGGSSNQQAIEDNTKRLMEQM...QTMQNAL..
Abalone_CLCC1    366  FLFMYFRYSVRLPFWLLSIEPGQREDNS...NVQLALQEQGKELK..........ALK..
Ictalurus_CLCC1  371  FIYSSSQAAVHHAFLR..PLRGGRQDPPPS.IAQQPAAPPIREREEHGEEVRYLLAG...
Anguilla_CLCC1   330  FMYGSAQAAIQHIIPR..PW.GGRQDPPPA.LVAQNPVPQLREAPPD......QPAG...

HUMAN_CLCC1      394  .GDADFHYRGQMGP..............TEQGPYAKTY.........EGRR.........
MOUSE_CLCC1      394  .GDADFSYRGPAGS..............IEQGPYDKMH.........ASKR.........
DANRE_CLCC1      423  ..DA.RQPLP.MRQ...DNRGNHVGNR.GDQGFRDANA.........PENR.........
XENLA_CLCC1      416  .GQLPRDN.DVVNI..............PKQQPLDDI.......................
OsHV1_ORF57      304  .G.LCYNKLFR.......................NKKY......................
AbHV1_ORF90      328  .S.LKIKK....................................................
IcHV1_ORF16A     310  NGVAAMEYKEA........SEETHGEG.EEEGPA..........................
AngHV1_ORF112    263  RGYLNRVAEFPA.................AERE...........................
Oyster_CLCC1     415  ...ESRETQF..T........ARMNDFER....LQKTAIEYSAAINVPDAP.MPTIPKEP
Abalone_CLCC1    411  ...EAHSSQM..N........EIMDVLKKDQQALEQQLANQETILSLPDVRRMVQVPPEK
Ictalurus_CLCC1  425  .GDANRAAQPRHHIRQEGNEANRMGNR.NSEEE...NR.........PEIR.........
Anguilla_CLCC1   377  .GDAPRQAVL................P.RRQGE...QR.........APVR.........

HUMAN_CLCC1      421  ...EILRERDVDLR.FQTGNKSPEVLRAFDV....PDAEAREHPTVVPSHKSPVLDTKPK
MOUSE_CLCC1      421  ...DALR.....QR.FHSGNKSPEVLRAFDL....PDTEAQEHPEVVPSHKSPIMNTNL.
DANRE_CLCC1      457  ...EEDRSMDIRQE.FSTKRTPVETLQATGN....TFPDDETDSQQRTQ..ELDSGANVE
XENLA_CLCC1      437  .................DGSNNPPVTAPAD............................PS
OsHV1_ORF57           ............................................................
AbHV1_ORF90           ............................................................
IcHV1_ORF16A          ............................................................
AngHV1_ORF112         ............................................................
Oyster_CLCC1     457  ...AVSRQRTTTPISQESGDLVPLTVKPHDTKSPIPCSEAELESSTEDS...........
Abalone_CLCC1    458  QGAEVMASRS.YPM.IAGGDRKPDLTGVEDQSNQVFCAAGGPEPQ...............
Ictalurus_CLCC1  462  ...QR...GPNRPR.IQQQRVYVETLRNADR....FYSGDETDTQQEAV..AEDLAQQV.
Anguilla_CLCC1   398  ...QR...RQNRRR.EDQAPVFVETLRQADH....PFSEDEIDAEHQED..HSEMEDHEL

HUMAN_CLCC1      473  ETGGILGEGTPK.ESSTESSQSAKPVSGQDTSGNT.....EGSPAAEKAQLKSE..AAGS
MOUSE_CLCC1      467  ETGELPGESTPT.E....YSQSAKDVSGQVPSA.G.....KSSPTVDKAQLKTD..SECS
DANRE_CLCC1      507  EE..VKVEEKEKKESFS...VDNKEQKE.TKSPDRSEPITSEPPSSIDVKTVGA..DQGN
XENLA_CLCC1      452  DTGQVKSNNTGE.PLV....QEDHSIKKSIKESRN.....DERPNTESPEA.........
OsHV1_ORF57           ............................................................
AbHV1_ORF90           ............................................................
IcHV1_ORF16A          ............................................................
AngHV1_ORF112         ............................................................
Oyster_CLCC1     503  ............ER......TINLPHGGQSPSRGRSSRLPLSKKNKTGEQLPT....QIS
Abalone_CLCC1    501  .............E......QLNI..SGDSEI..KNVQLALQEQGKELKALKEAHSSQMN
Ictalurus_CLCC1  508  .......E....EEPLT...LSNS.PKEEHDSSASNTEMKHTQPASKEHSAKSD..TKDN
Anguilla_CLCC1   445  PTGGALLEEE..EENVV...TENENLSGDSQCNSNKDRLKSENPSG..HSKSAQ..QKGE

HUMAN_CLCC1      525  PDQGSTYSPARGVAG..................PRGQD...PVSSPCG............
MOUSE_CLCC1      514  PPGGCPPSK.EAAVA..................AHGTE...PVSSPCG............
DANRE_CLCC1      559  EHLMCTKRKWAAQNGFKLQVILCE.INSEASADLPEEEECFSFKHPVQ...ETQS.....
XENLA_CLCC1      493  ....KPQRP...........................EE...PVVETLR...ST.......
OsHV1_ORF57           ............................................................
AbHV1_ORF90           ............................................................
IcHV1_ORF16A          ............................................................
AngHV1_ORF112         ............................................................
Oyster_CLCC1     541  SAIDSS..........HTHVISPETVLSQ....ETSSR......................
Abalone_CLCC1    538  KIMDVLKK...DPQALEQQLANQETILSL.....PDVRRMVQVPPEKQGAEVMASRSYPM
Ictalurus_CLCC1  551  KTSGNTERN.AVGDS...........ASVGKAASKKEDRVENIGTPVQ...ETPVYIYST
Anguilla_CLCC1   496  DPSRNSQRP.RVPDG...........LSEEKTTSVTQERVENIGSPIQ...ETAFQS...

HUMAN_CLCC1           ........................................
MOUSE_CLCC1           ........................................
DANRE_CLCC1           ........................................
XENLA_CLCC1           ........................................
OsHV1_ORF57           ........................................
AbHV1_ORF90           ........................................
IcHV1_ORF16A          ........................................
AngHV1_ORF112         ........................................
Oyster_CLCC1          ........................................
Abalone_CLCC1    590  IAGGDRKPDLTGVEDQSNQVICAAGGPEPQEQLNISGDSR
Ictalurus_CLCC1  596  SH.....SVL..QSTQYCTALCS.................
Anguilla_CLCC1        ........................................
```

**Supplementary Figure 2. Sequence alignment of 8 cellular and 4 herpesviral CLCC1 homologs.** Multiple sequence alignment of CLCC1 homologs from *Homo sapiens* (Human_CLCC1), *Mus musculus* (MOUSE_CLCC1), *Danio rerio* (zebrafish; DANRE_CLCC1), *Xenopus tropicalis* (western clawed frog; XENLA_CLCC1), osterid herpesvirus 1 (OsHV1_ORF57), abalone herpesvirus 1 (AbHV1_ORF90), Ictalurid herpesvirus 1 (IcHV1_ORF16a), anguillid herpesvirus (AngHV1_ORF112), *Crassostrea gigas* (Pacific oyster; Oyster_CLCC1), *Haliotis rubra* (blacklip ablone; Ablone_CLCC1), *Ictalurus punctatus* (channel catfish; Ictalurus_CLCC1), and *Anguilla rostrata* (American eel; Anguilla_CLCC1). Similar residues are highlighted in yellow. Conserved residues are highlighted in red. Sequence alignment was generated using Clustal Omega[1] and rendered using ESPript 3.0[2] (https://espript.ibcp.fr).

**a**

| Gene.ID | log(FC) | -log(p) | Description |
|---|---|---|---|
| CLCC1 | 4.133 | 9.266 | Chloride channel CLIC-like protein 1 |
| CCNC | 0.6536 | 4.171 | Cyclin-C |
| NIPAL1 | 0.7891 | 3.821 | Magnesium transporter NIPA3 |
| TP53BP1 | 0.4226 | 3.525 | TP53-binding protein 1 |
| WDR20 | 0.4626 | 3.438 | WD repeat-containing protein 20 |
| BTAF1 | 0.9582 | 3.344 | TATA-binding protein-associated factor 172 |
| DLG4 | 0.3615 | 3.145 | Disks large homolog 4 |
| MOV10L1 | 0.485 | 3.134 | RNA helicase Mov10l1 |
| VCPKMT | 0.6487 | 3.012 | Protein-lysine methyltransferase METTL21D |

**b**

| Gene.ID | log(FC) | -log(p) | Description |
|---|---|---|---|
| CSNK1E/TPTEP2-CSNK1E | -0.5074 | 4.168 | Casein kinase I isoform epsilon |
| CRTAP | -0.9859 | 4.11 | Cartilage-associated protein |
| EMD | -0.8608 | 4.026 | Emerin |
| CBWD6 | -0.862 | 3.997 | COBW domain containing 6 |
| ZNF7 | -0.7699 | 3.961 | Zinc finger protein 7 |
| IL6ST | -1.413 | 3.95 | Interleukin-6 receptor subunit beta |
| SEC22B2P/SEC22B3P | -0.8272 | 3.895 | Vesicle-trafficking protein SEC22b homolog B2/B3 |
| SBNO2 | -0.8282 | 3.848 | Protein strawberry notch homolog 2 |
| MUC16 | -0.4844 | 3.764 | Mucin-16 |
| NOMO2/NOMO3 | -0.6829 | 3.559 | Nodal modulator 2/3 |
| F8A1/F8A2/F8A3 | -0.7068 | 3.423 | 40-kDa Huntingtin-associated protein |
| PCSK1 | -0.8288 | 3.423 | Neuroendocrine convertase 1 |
| ABCE1 | -0.8815 | 3.415 | ATP-binding cassette sub-family E member 1 |
| H3C14/H3C15 | -0.9308 | 3.41 | H3 clustered histone 14/15 |
| U2AF1/U2AF1L5 | -0.596 | 3.364 | Splicing factor U2AF 35 kDa subunit/U2AF1 like 15 |
| PHF20L1 | -0.4748 | 3.354 | PHD finger protein 20 like 1 |
| NDUFS5 | -0.7555 | 3.321 | NADH dehydrogenase [ubiquinone] iron-sulfur protein 5 |
| POTEE | -0.9869 | 3.301 | POTE ankyrin domain family member E |
| NPIPB7 | -0.8821 | 3.291 | Nuclear pore complex interacting protein family member B7 |
| CBWD3 | -0.9364 | 3.278 | COBW domain containing 3 |
| SPAG1 | -0.8011 | 3.263 | Sperm-associated antigen 1 |
| OR2AG2 | -0.4425 | 3.262 | Olfactory receptor 2AG2 |
| UQCRB | -0.7618 | 3.202 | Cytochrome b-c1 complex subunit 7 |
| LSM10 | -1.205 | 3.171 | U7 snRNA-associated Sm-like protein LSm10 |
| ANAPC1 | -0.6037 | 3.151 | Anaphase-promoting complex subunit 1 |
| CLUAP1 | -0.9972 | 3.147 | Clusterin-associated protein 1 |
| CYTH1 | -1.296 | 3.134 | Cytohesin-1 |
| FGF3 | -0.7526 | 3.126 | Fibroblast growth factor 3 |
| MASP2 | -0.8831 | 3.061 | Mannan-binding lectin serine protease 2 A chain |
| NEK9 | -0.6475 | 3.029 | Serine/threonine-protein kinase Nek9 |
| SAA1 | -1.171 | 3.015 | Serum amyloid protein A(2-102) |
| PIKFYVE | -1.607 | 3.003 | 1-phosphatidylinositol 3-phosphate 5-kinase |

**Supplementary Table 1. Genome-wide CRISPR screen high confidence candidates. a)** 9 high-confidence candidate positive regulators (decreased nuclear egress when the gene is depleted), log(FC) > 0, p-value < 0.001. Top hit, CLCC1, is labelled in red. **b)** 32 high-confidence candidate negative regulators (increased nuclear egress when the gene is depleted), log(FC)<0, p-value < 0.001. EMD is labelled in green.

| Gene.ID | log(FC) | -log(p) |
|---|---|---|
| EMD[3] | -0.8608 | 4.026 |
| LMNA[4] | -1.06 | 1.813 |
| SLC35E1[5] | -0.7265 | 1.678 |
| CHMP4C[6] | 0.5457 | 1.602 |
| PRKCB[7] | 0.2192 | 1.425 |
| PRKD2[8] | -0.6528 | 1.406 |
| PRKCG[7] | -0.5937 | 1.044 |
| PRKCA[7] | -0.2916 | 0.9876 |
| PRKD1[8] | -0.3519 | 0.924 |
| CHMP4B[6] | -0.5279 | 0.8362 |
| TOR1A[9] | 0.3163 | 0.6801 |
| SLC3A2[10] | -0.3851 | 0.6346 |
| DDX3X[11] | -0.3417 | 0.595 |
| ITGB1[10] | -0.4896 | 0.5739 |
| CHMP4A[6] | -0.2494 | 0.3303 |
| TSG101[12] | -0.2126 | 0.2406 |
| C1QBP(p32)[13] | -0.1927 | 0.2246 |
| PDCD6IP (ALIX)[6] | -0.1532 | 0.1751 |
| VAPB[14] | -0.06719 | 0.1154 |
| SUN2[15] | 0.05463 | 0.08297 |
| TOR1B[9] | -0.01995 | 0.02847 |
| TOR1AIP2[16] | 0.01504 | 0.01238 |

**Supplementary Table 2. Host factors previously reported as contributing to HSV-1 nuclear egress.**

| Family | Protein Description | Accession No. |
|---|---|---|
| *Malacoherpesviridae* | ORF57 [Ostreid herpesvirus 1] | ASK05584.1 |
| *Malacoherpesviridae* | ORF56 [Chlamys acute necrobiotic virus] | ADD24788.1 |
| *Malacoherpesviridae* | ORF100 [Malaco herpesvirus 1] | DBA11801.1 |
| *Malacoherpesviridae* | ORF90 [Abalone herpesvirus 1] | YP006908742.1 |
| *Alloherpesviridae* | ORF16A [Ictalurid herpesvirus 1] | QAB08501.1 |
| *Alloherpesviridae* | ORF112 [Anguillid herpesvirus 1] | APD76276.1 |
| *Alloherpesviridae* | ORF94 [black bullhead herpesvirus] | YP_009447842.1 |
| *Alloherpesviridae* | ORF15 [Silurid herpesvirus 1] | AVP72192.1 |

**Supplementary Figure 3. CLCC1 homologs encoded in herpesviral genomes.** A list of herpesviral homologs of CLCC1 and their accession numbers.
